# The DNA-to-cytoplasm ratio broadly activates zygotic gene expression in Xenopus

**DOI:** 10.1101/2021.04.18.440334

**Authors:** David Jukam, Rishabh R Kapoor, Aaron F Straight, Jan M. Skotheim

## Abstract

**Summary:** In multicellular animals, the first major event after fertilization is the switch from maternal to zygotic control of development. During this transition, zygotic gene transcription is broadly activated in an otherwise quiescent genome in a process known as zygotic genome activation (ZGA). In fast developing embryos, ZGA often overlaps with the slowing of initially synchronous cell divisions at the mid-blastula transition (MBT). Initial studies of the MBT led to the nuclear-to-cytoplasmic ratio model where MBT timing is regulated by the exponentially increasing amounts of some nuclear component ‘N’ titrated against a fixed cytoplasmic component ‘C’. However, more recent experiments have been interpreted to suggest that ZGA is independent of the N/C ratio. To determine the role of the N/C ratio in ZGA, we generated *Xenopus* frog embryos with ∼3-fold differences in genomic DNA (*i.e.*, “N”) by using *X. tropicalis* sperm to fertilize *X. laevis* eggs with or without their maternal genome. Resulting embryos have otherwise identical *X. tropicalis* genome template amounts, embryo sizes, and *X. laevis* maternal environments. We used the *X. tropicalis* paternally derived mRNA to identify a high confidence set of exclusively zygotic transcripts. Both ZGA and the increase in cell cycle duration are delayed in embryos with ∼3-fold less DNA per cell. Thus, DNA is an important component of the N/C ratio, which is indeed a critical regulator of zygotic genome activation in *Xenopus* embryos.

## Introduction

Animal development is initially driven by the large pool of maternal factors until the maternal-to-zygotic transition, when the developing embryo initiates transcription to begin to take control of its own development ^1^. The timing of embryonic gene activation requires coordination of maternal mRNA translation and chromatin changes with cell cleavage divisions that increase cell number to prepare for gastrulation ^2^. Yet our understanding for how the zygotic genome initiates transcription in the hours after fertilization remains incomplete.

Classic work in rapidly developing vertebrates produced a model where changes in the nucleo-to-cytoplasmic ratio (N/C) induced simultaneous events at a mid-blastula transition (MBT) comprising cell cycle lengthening and the activation of the zygotic genome ^3, 4^. The N/C ratio model proposed that the rapid cell divisions of early development that proceed with negligible cell growth change the ratio of one or more nuclear components relative to cytoplasmic volume to induce the MBT. The N/C model was initially based on experiments manipulating DNA content or mechanically separating cytoplasm and then assaying their effects on cell cycle durations. In multiple species, increasing ploidy or decreasing cytoplasm advanced the MBT, and decreasing ploidy delayed the MBT ^3, 5–8^.

While the classic N/C ratio likely controls cell cycle changes, it has been unclear whether transcription was similarly N/C ratio-dependent ^9^. It is now apparent in several fast developing species that hundreds of genes are actively transcribed during cleavage divisions before the MBT, indicating that zygotic genome activation does not necessarily occur at the same time as other MBT events ^10–18^. Different genes are activated at different times and are regulated by multiple maternally provided transcriptional activators ^19–22^. Moreover, recent work has called into question the extent to which the N/C ratio influences transcription initiation in fish and frogs ^21, 23^. Although there exists evidence that MBT cell cycle lengthening permissively allows transcription of long genes in *Drosophila* ^15, 24^, it has been suggested that neither the N/C ratio or cell cycle slowing affects large-scale <u>z</u>ygotic <u>g</u>enome <u>a</u>ctivation (ZGA) timing in *Xenopus* frogs ^14^. Instead, models suggest that the timing of zygotic gene expression is largely due to activator concentration changes, histone acetylation near promoters, and chromatin state changes ^14, 23, 25^, and may therefore be unrelated to the N/C-ratio. One possibility to bridge this N/C ratio divide would be if a subset of genes were N/C-ratio regulated while others were not, as was suggested to be the case in *Drosophila* ^26^. Thus, the extent to which the N/C ratio influences transcriptional timing remains ambiguous.

Here, we sought to determine the extent to which the N/C ratio regulates zygotic genome activation by examining frog embryos. Prior experiments in *Xenopus* frogs supporting N/C ratio control of ZGA were limited to assaying either exogenously injected reporter constructs, RNA Pol III mediated transcripts, or a handful of *in vivo* transcribed mRNA genes ^4, 27–29^. Furthermore, interpretations of prior studies in vertebrate embryos that underwent altered ploidy in haploids are confounded because any changes in ploidy also change the DNA template for gene expression. To overcome limitations of previous studies, we exploited the viability of *Xenopus laevis* and *Xenopus tropicalis* hybrids to create embryos wherein DNA content can be manipulated *in vivo*, while maintaining a constant genomic template for RNA. We compare the zygotic transcriptome in embryos with a ∼3-fold difference in genomic DNA and find that timing for nearly all ZGA genes at the MBT is regulated by the DNA-to-cytoplasm ratio. Increased transcription at the MBT was due to an increasing number of transcripts per genome in addition to the exponentially increasing number of genomes. Our results, combined with recently reported cell size effects – wherein a critical cell size threshold may be necessary for bulk transcriptome activation in individual embryonic cells in *Xenopus* ^30^ – support the view that DNA is a critical parameter for “N”, and that cell size (or rather, cytoplasmic volume) is the critical parameter for “C”. We suggest a model wherein activators and chromatin changes initiate gene expression, but expression levels and timing for ZGA genes are additionally regulated by the N/C-ratio at the MBT.

## Results

### Using *X. laevis* and *X. tropicalis* hybrids to investigate ZGA

To assess whether the N/C ratio regulates zygotic gene expression in Xenopus embryos, we focused on the “N” numerator that relates to some parameter(s) of the nucleus. Prior work suggests that one important component of “N” is likely DNA or chromatin, as increased or decreased ploidy manipulations cause an advance or delay in cell cycle lengthening, respectively. The constant length of DNA per cell can then serve as a yardstick to “measure” the exponentially decreasing amount of cytoplasm per cell, as proposed in titration models for MBT induction ^4^.

To test whether a change in the amount of DNA per cell affects transcription, we required an experimental approach in which total embryonic DNA content can be manipulated without changing the maternal cytoplasm or amount of DNA template for gene expression. Prior experiments linking ploidy to MBT transcriptional timing relied on injection of exogenous plasmid DNA ^4^ or haploid embryos ^7, 29^. However, changing the amount of DNA template for transcription confounds interpretation because one cannot distinguish whether expression level changes are due to changes in timing or due to gene dosage.

To circumvent the limitations of previous experiments to determine whether or what portion of zygotic gene expression is regulated by the N/C ratio, we took advantage of the ability of related *Xenopus* frog species to cross-fertilize and form hybrid embryos that are competent to develop through embryogenesis and morphogenesis into adult frogs ^31–33^. *X. laevis* is allotetraploid (∼2.8 x 10^9^ Gb DNA; 2n=36 chr) and *X. tropicalis* is diploid (∼1.4 x 10^9^ Gb; 2n=20 chr) ^34, 35^. Thus, fertilizing *X. laevis* eggs with *X. tropicalis* sperm results in hybrids whose cells retain one copy of each species’ genome – the *Xenopus* short (S) and long (L) subgenomes and a haploid *tropicalis* genome – for 3 haploid genome equivalents (∼4.2 x 10^9^ Gb; N=28 chr). This chromosome complement is stable throughout development.

Using *X. laevis* (egg) x *X. tropicalis* (sperm) hybrids as a baseline, we generated embryos with ∼3-fold less DNA content by irradiating the *X. laevis* egg before fertilization with *X. tropicalis* sperm (Figure 1A). A short pulse of a ∼350 nm UV laser was sufficient to crosslink the *X. laevis* egg genome without otherwise damaging the embryos or the ability of these embryos to be fertilized by wild-type *X. tropicalis* sperm (Figure 1C-D) ^32, 36^. Crosslinking the maternal genome results in rapid extrusion or loss of the entire maternal genome in the embryo such that only the haploid paternal *X. tropicalis* genome remains ^36^. We refer to these embryos as nucleocytoplasmic hybrids (hereafter, “cybrids”) since they contain the maternal *X. laevis* cytoplasm, and the *X. tropicalis* nucleus and DNA. We performed karyotyping at the neurulation and tailbud stages to measure single-cell ploidy for both hybrids and cybrids. Hybrids contain the expected 28 chromosomes (18 from *laevis* and 10 from *tropicalis*), whereas the cybrids contained only 10 chromosomes (Figure 1C-D).

**Figure 1.**
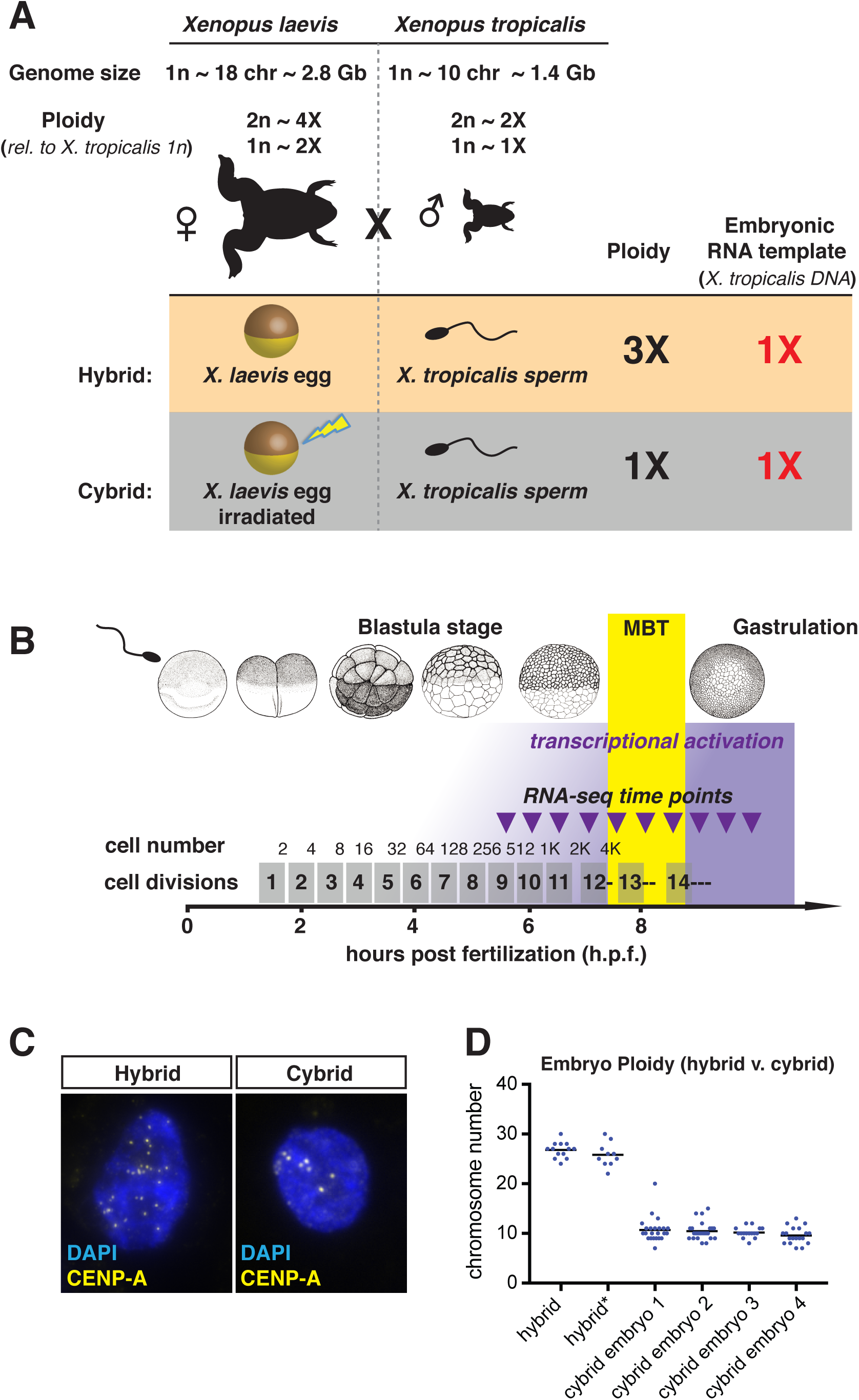
*Xenopus* hybrid and cybrid embryos have a 3-fold difference in DNA per cell. (A) Schematic illustrating the experimental design. *Xenopus laevis* has ∼2X more DNA than *Xenopus tropicalis*. *X. laevis* eggs fertilized with *X. tropicalis* sperm generate hybrid embryos. UV-irradiation destroys the genome in *X. laevis* eggs so that subsequent fertilization results in cybrid (cytoplasmic hybrid) embryos containing 3-fold less DNA per cell than hybrid embryos. The *X. laevis* maternal environment and the *X. tropicalis* genome template for RNA expression are otherwise identical in both conditions. (B) Schematic depicting the developmental timeline of early *Xenopus* embryo development. Time after fertilization (hours) is on the bottom axis. Cleavage cell division number is above in grey rectangles. Embryo cell number following each synchronous division is above-right of the grey rectangles. The MBT (yellow bar) occurs at the ∼12th cell division (4000-cell stage), which corresponds to the 8.5 Nieuwkoop and Faber (NF) morphological embryonic stage. Dashed lines after the 12th, 13th, and 14th divisions indicate loss of cell division synchrony. Purple triangles indicate time of embryo collection for the transcriptomic time-series that begins at 5.5 hours post-fertilization. (C) Images of representative interphase nuclei from hybrid (left) and cybrid (right) embryos, stained to visualize DNA (DAPI, blue) and centromeres using primary antibodies against the centromere-specific protein CENP-A (yellow). The number of interphase centromeres is equivalent to the number of chromosomes. (D) Graph displaying the chromosome counts in nuclei examined from hybrids and cybrids. Each data point represents a single nucleus, and each cluster is from a different embryo. The number of chromosomes decreases from ∼28 in hybrids to ∼10 in cybrids. The single nucleus (arrow) with 20 chromosomes in cybrid-embryo-1 was noted to be at the mitotic stage with the expected double *X. tropicalis* chromosome complement.

We observed normal development in hybrid embryos from the blastomere stage across gastrulation and neurulation, as previously reported (Gibeaux et al., 2018b). Swimming tadpoles were more morphologically similar to wild-type *X. laevis* tadpoles than were *X. laevis* haploids, consistent with prior work ^32^. Therefore, *X. laevis* (egg) x *X. tropicalis* (sperm) hybrids appear developmentally “normal” during embryogenesis in contrast to haploids in either species which are smaller and have a greater number of morphological aberrations visible at the tailbud stage ^32^ (data not shown). Like wild-type and hybrid embryos, cybrid embryos were fertilized with high efficiency and displayed synchronous cleavage divisions from the 1-cell stage until the MBT. Cybrid embryos also develop normally through the MBT and early gastrulation (eventually dying post neurulation) and are thus suitable for studying early embryonic gene expression ^32^. The resulting hybrid and cybrid embryos therefore provide a ∼3-fold difference in genomic DNA while other aspects of embryo development are maintained.

### Defining *X. tropicalis* ZGA genes

The first step in determining the effect of DNA content on zygotic transcription is to determine the set of genes that are transcriptionally activated during the MZT. While defining ZGA genes is typically difficult due to the large maternal pool of mRNA, it is straightforward to do with hybrids because many transcripts from the *X. tropicalis* genome can be distinguished from the *X. laevis* maternal pool. We therefore assayed gene expression in hybrids across the MBT every 30 minutes, a time equivalent to ∼1 cleavage cell division, beginning at 5 hours post-fertilization (hpf) using RNA-seq of the non-ribosomal transcriptome. The transcriptomic time series data from up to 11 independent samples across 3 replicates was of high quality and reproducible with high correlations between neighboring time points and replicates at each stage (Figure S1A-B). Because the transcriptome composition across early embryogenesis is extremely dynamic compared to somatic cells, we added exogenous RNAs during embryo collection to normalize our samples. Spike-ins showed reproducibility across replicates and time points, and matched their expected abundance (Figure S2A-C). This normalization approach using spike-ins allows us to perform rigorous quantitative analysis on the resulting data.

*X. tropicalis* zygotic gene expression in hybrids increases gradually through the first few time points and then more extensively at the MBT (Figure S3A). Furthermore, we could accurately distinguish most *X. tropicalis* zygotic transcripts from their orthologous maternal *X. laevis* S and L transcripts despite their high abundance (*e.g., cdk9, srsf6, vegt, sox2, pou5f3.3, and ets2*) (Figure S3B-D). The vast majority (∼99%) of sequences from early developmental stages before 7 h.p.f. were *X*. *laevis* maternal genes, as expected (Figure S3A). A subpopulation of *X. tropicalis* genes that exhibited high early expression were likely *X. laevis* maternal genes mis-attributed to *X. tropicalis* due to sequence similarity. Our filter excluded these genes from further analysis. This approach allowed us to identify a set of 595 zygotically expressed *X. tropicalis* genes in hybrids that we define as “ZGA-genes”, which all satisfy specific threshold criteria (see methods) (Figure 2A-B). The expression of the remaining transcriptome, composed primarily of *X. laevis* maternal genes and *X. tropicalis* genes that did not pass our filters, remained broadly unchanged through the MBT (5.5 hpf compared to 9.5 hpf) (Figure 2B). Although the ZGA set did not include the first expressed transcript in *Xenopus*, *miR-427*, due to extremely high early expression, as expected, this miRNA exhibited a clear ZGA-like trend and was 128-fold more expressed post-MBT vs pre-MBT (Figure 2B). In addition, our defined ZGA genes are overrepresented for processes such as transcription, gastrulation, and germ layer formation GO terms, as we would expect from bona fide *Xenopus* ZGA genes (Figure S4A).

**Figure 2.**
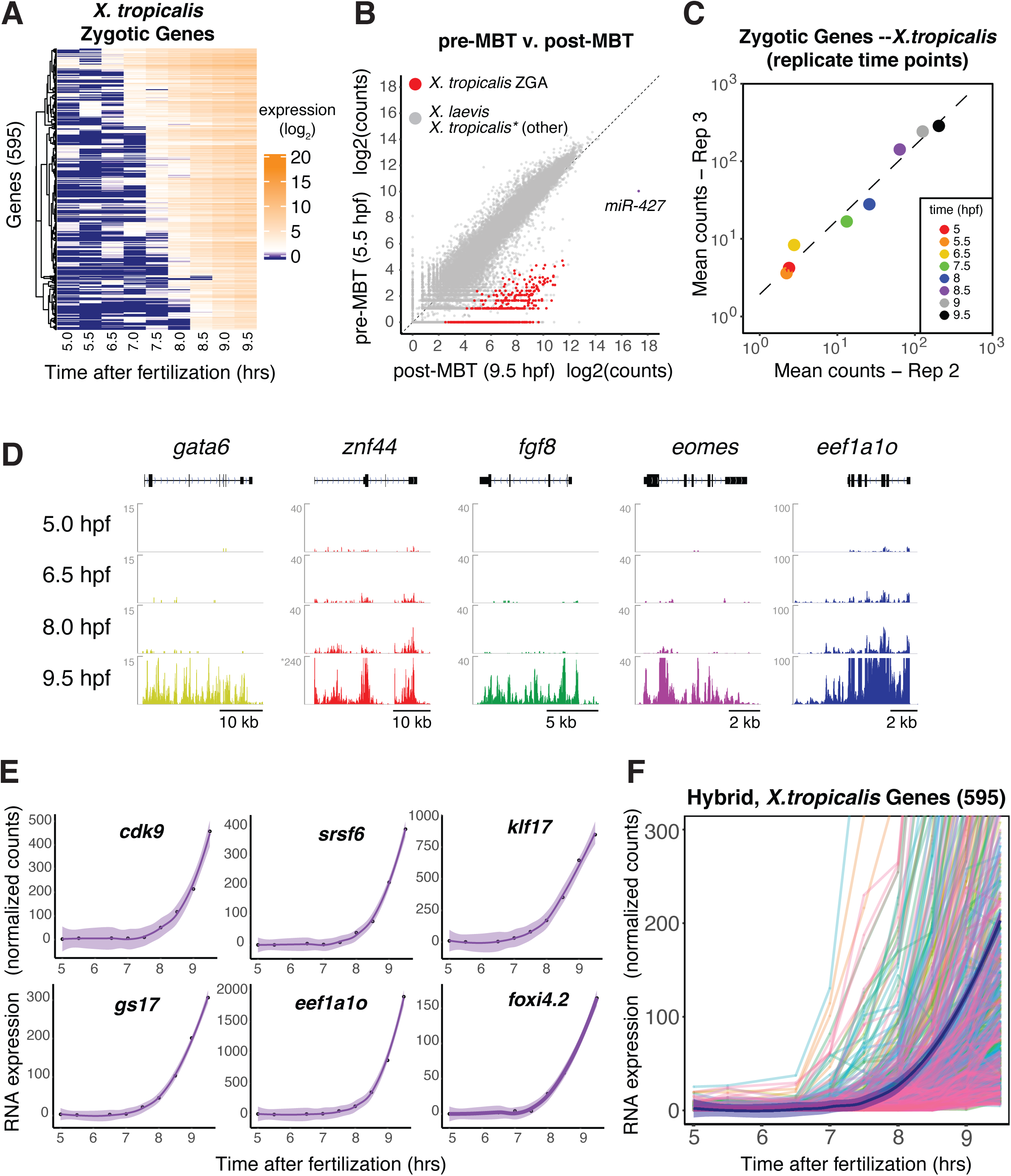
Identification of zygotically expressed *X. tropicalis* genes in hybrids. (A) Heatmap of defined zygotic *X. tropicalis* gene expression in hybrids across the MBT. Genes were hierarchically clustered across time points by their normalized expression. Expression counts were log2 normalized prior to clustering. Color value scale represents min (expression = 0, dark blue), mean (exp=10), and max (exp=20, dark orange) expression. All expression values in all figures are ERCC-spike-in normalized gene counts prior to additional normalization unless otherwise noted. (B) Scatterplot showing gene expression for *X. tropicalis* ZGA genes (red) and remaining genes (gray). Log2-normalized counts expression values are compared before and after the MBT, at 5.5 (Y-axis) and 9.5 (X-axis) hpf, respectively. Each point represents a single gene. The first transcript expressed in *X. tropicalis*, *miR-427*, is annotated (purple). Dashed diagonal line is X=Y. (C) Scatterplot of the gene counts centroids (mean X and mean Y) from each time point for the 595 ZGA genes, for all common time points from two biological replicates. Each time point is represented by a separate color. Axes display log2-normalized counts. Dashed line is a linear fit to the data. (D) Genome browser images showing gene expression signal (spike-in normalized read counts) for zygotic *X. tropicalis* transcripts for 5 representative ZGA genes--*gata6*, *znf44*, *fgf8*, *eomes*, and *eef1a1o*. Signal tracks include merged data from replicates 2 and 3. Read count shown in Y-axis. Signal intensity was originally at base-pair resolution (unbinned) with mean signal over windows shown for clarity. Transcript structure at top in black (boxes = exons; line = introns). Note the presence of intronic signal in zygotic *X. tropicalis* genes (black arrows) that likely represents nascent transcription. Grey asterisk indicates where the 9.5 hpf Y-axis for *znf44* is scaled 6-fold higher to better display the complete intron/exon signal. Note that the 9.5 hpf Y-axis for *eef1a1o* is scaled to match the earlier time points, and therefore does not display the entire gene signal. (E) Expression time-course profiles for individual ZGA genes. Gene expression is normalized using spike-in counts. Points are overlaid with a lowess fit (purple line) and standard error of the fit (purple shading). (F) Time-course profiles for each defined ZGA gene from hybrids. A composite lowess fit (thick black line) and standard error (purple shading) from all ZGA genes is overlaid. Individual colors represent a single gene. (N=595)

The overall trend for our defined ZGA set is low or zero expression before the MBT, which is followed by an hour of gradual increase, and then a sharp increase at the MBT (Figure 2C-F) ^16, 17^. This trend is consistent across replicates from independent clutches as the replicates show high reproducibility at all common time points (Figure 2C). Our defined zygotic gene set includes many canonical ZGA genes such as *gata6*, *fgf8, eomes, cdk9, klf17, and gs17* (Figure 2D-E) ^17^, as well as gene families that regulate gastrulation and germ layer development (Figure S4A). In addition, the ZGA gene set shows statistically significant overlap with previously identified *X. tropicalis* zygotic genes (Figure S4B) ^16, 17^. Our RNA-seq approach is sensitive enough to detect relatively lowly expressed zygotic genes (Figure 2E-F), and the maximal expression of each gene ranges over 1000-fold. We conclude that our approach to define *X. tropicalis* ZGA genes in hybrids is sensitive and accurate, and provides a high-confidence ZGA gene set for rigorous analysis of gene expression timing.

### Greater DNA broadly induces earlier ZGA timing

Having defined a high confidence set of *X. tropicalis* ZGA genes, we sought to determine whether their transcription was sensitive to cellular DNA content. The DNA-to-cytoplasm ratio model predicts that altering DNA content in cells of otherwise similar size will cause a shift in the onset of zygotic gene expression around the time the MBT initiates at the 12th-13th cleavage divisions in *Xenopus* ^3, 37^.

To test whether a decrease in per-cell DNA content can delay the ZGA, we compared high-resolution RNA-seq time series expression profiles from hybrids to that of cybrids, which contain ∼3-fold less DNA per cell. Hybrid and cybrid embryos from each biological replicate were sampled in matched time points from the same clutch and fertilization event. Two individual samples, and their matching time points, were removed due to low quality from 2 replicates. This resulted in 3 gene expression time-courses with matched hybrids and cybrids across the MBT. Cybrids developed normally through the MBT and early gastrulation ^32^. We found that the population of maternal *X. laevis* mRNAs in cybrids was both highly and stably expressed across the cleavage stages, and was very similar to the maternal mRNA pool in hybrids including mRNAs for several transcription factors critical for genome activation (Figure S3E) ^21^.

A comparison of gene expression during early development in hybrids and cybrids was in broad agreement with the DNA-to-cytoplasm ratio model. We found that ZGA genes were either not expressed or expressed at low levels at early time points, and not noticeably different in hybrids and cybrids (Figure 3A). At the MBT, however, the ZGA genes had decreased expression as a group in the cybrids relative to the hybrids, suggesting a delay in transcriptional activation in the cybrids that have 3X less DNA per cell (Figure 3B). After the MBT at 9.5 hpf, ZGA genes were highly and similarly expressed in hybrids and cybrids, suggesting that expression levels for delayed genes in the cybrid eventually recovers to that of the hybrid (Figure 3C). Moreover, this cybrid ZGA expression level recovery to that of hybrids indicates that the observed expression delay is not due to grossly aberrant or sick embryos. Together, these trends reveal that the expression of ZGA genes is reduced in embryos with less DNA, consistent with the N/C-ratio influencing ZGA timing directly through genomic DNA.

**Figure 3.**
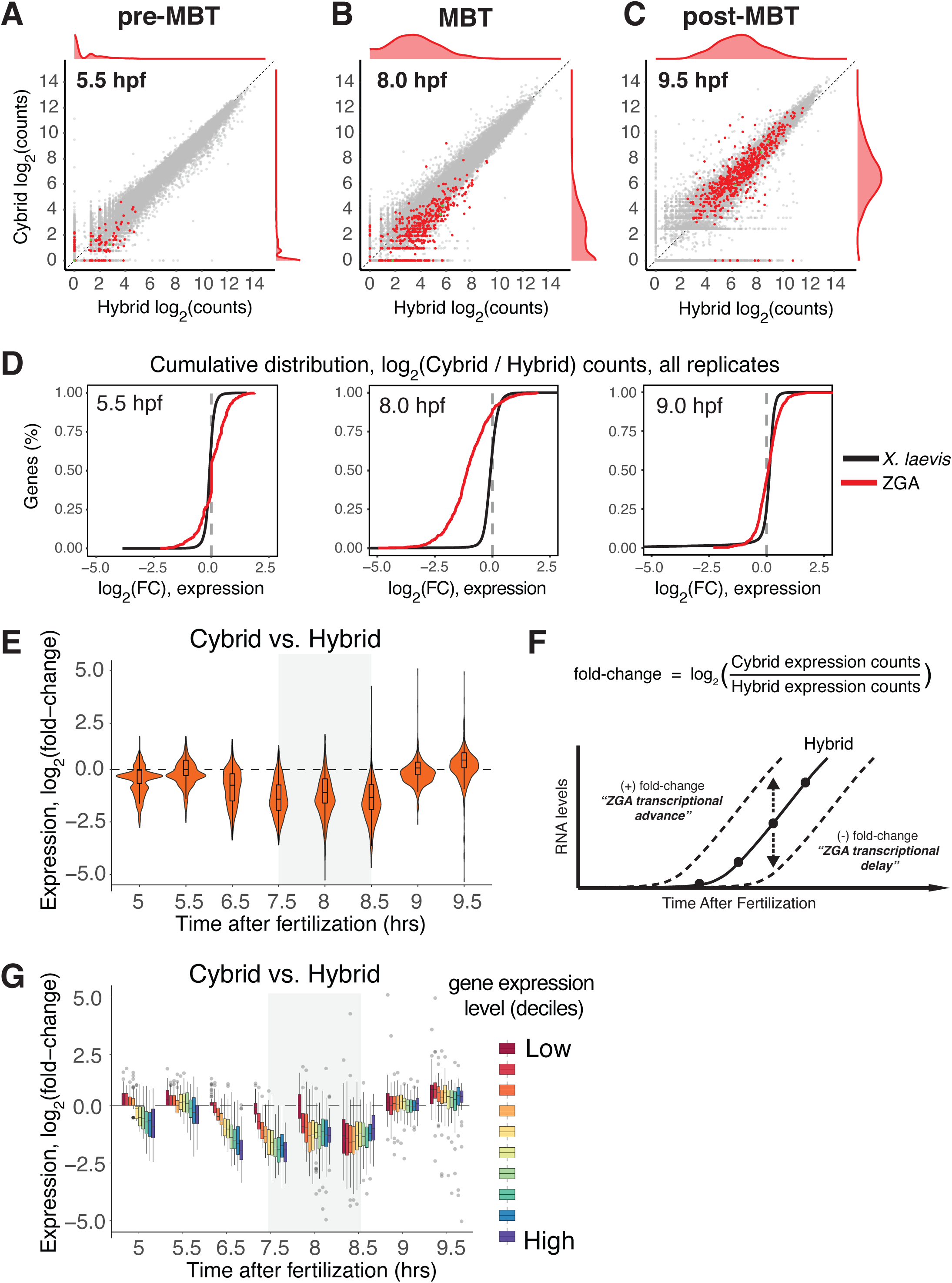
Embryos with less DNA have less zygotic gene expression specifically at the MBT. (A) Scatterplot of gene expression values for all genes before the MBT at 5.5 hours post fertilization (hpf). In (A-C): ZGA genes are shown in red, while all other genes are shown in grey; marginal distribution of ZGA gene numbers shown on right and top; X- and Y-axes show log2(counts) of spike-in normalized RNA-seq gene counts. (B) Hybrid vs cybrid gene expression values at the MBT, at 8.0 hpf. (C) Hybrid vs cybrid gene expression values after the MBT, at 9.5 hpf. (D) Cumulative distribution of log2(Cybrid – Hybrid) fold-change values of each ZGA gene’s expression, before the MBT (5.5 hpf), at the MBT (8.0), and after the MBT (9.0 hpf). A negative value is consistent with delayed expression in Cybrids as compared to Hybrids. Red line is defined *X. tropicalis* ZGA genes only; blue line is all *X. laevis* (two-sided Kolmogorov-Smirnov test: D=0.32 (5.5 hpf); D=0.69 (8.0 hpf); D=0.22 (9.5 hpf); p-value < 10^-10^ for all). (N=595) (E) Violin plot displaying the distribution of fold-changes for each ZGA gene’s expression in cybrid relative to hybrid embryos. Box inside shows mean and 1st and 3rd quartiles. Hybrid embryos are the reference, so a decreased fold-change indicates decreased expression in cybrids. Fold-changes are population means of every ZGA gene’s log2-normalized fold-change mean from 3 replicates. Each time point contains 2 or 3 replicates. (N=595) (F) Schematic of fold-change approach to compare transcriptomic time-series data. (G) Distribution of log2-normalized fold-changes of each ZGA gene’s expression in cybrid vs hybrid embryos, with genes binned according to expression level deciles. Expression level is per gene mean across all time points. Highest expression decile in purple, lowest in dark red. (N=595)

Having seen that higher DNA content broadly increases gene expression, we next sought to analyze how DNA content regulated the timing of transcriptional activation of individual genes. To do this, we first analyzed how individual gene expression levels differ at each time point in the cybrid relative to the hybrid (Figure 3F). During the cleavage stages before the MBT (*e.g.*, 5.5 hpf) as well as after the MBT (*e.g.*, 9.0 hpf), the distribution of per-gene fold-changes was centered around zero in both ploidy conditions (Figure 3D). In contrast, at the MBT (8.0 hpf), the majority of ZGA genes exhibited decreased expression levels in cybrids (Figure 3D, middle), which is consistent with more DNA advancing zygotic expression. This trend is also present when examining the mean fold-change across all time points in all replicates (Figure 3E, S5A-C). The initial delay in gene expression is slightly dependent on the individual gene’s maximum expression level (Figure 3G), likely due to the technical issue of more highly expressed genes more rapidly reaching our detection threshold after initiating transcription. However, this relationship is absent during the MBT (*e.g*., 8.0 and 8.5 hpf) (Figure 3G). The distribution of expression differences for hybrids and cybrids in the ZGA gene set roughly followed a gaussian distribution with no clearly demarcated subpopulations of genes with unchanged or increased expression (Figure S5D). The lack of obvious subpopulations within the ZGA set suggests that the majority of ZGA genes are delayed in embryos with less DNA per cell rather than there being a clearly defined subset of genes expressed independently of the N/C-ratio, as has been reported in *Drosophila* ^26^.

Taken together, our analysis so far shows that the vast majority of zygotically expressed *X. tropicalis* genes are delayed in embryos with less DNA per cell, consistent with an influence of the N/C-ratio on ZGA timing. As an independent approach to compare ZGA dynamics in hybrids and cybrids, we assessed when a specific gene sharply increases its expression in each condition. Rather than manually annotate regions of expression increase or choosing a simple threshold to determine activation times, we instead developed an algorithm that could be applied in a uniform manner to all genes without bias. To do this, we first normalized the gene expression time series in both hybrids and cybrids, and applied a filter to systematically remove outlying data points or genes with highly aberrant expression curves (Figure S6A), resulting in 547 ZGA genes from our original set of 595. We then estimated continuous gene expression levels across the time course using smoothing splines with a uniform smoothing parameter (Figure S6B and 4A) ^38^. This algorithm was applied to both the hybrid and cybrid time series for the two replicates containing eight or more time points. The “activation time” (*t*_Act_) for each gene was determined by calculating 10% of the maximum expression along the profile. This approach was superior to fitting the time series gene expression data with alternative models, such as hinge functions or exponential functions (see methods for discussion), and was consistent across replicates (Figure S7A-B). We note that *t*_Act_ is unlikely to describe the initial transcription of each zygotic mRNA, which occurs up to several hours before the MBT for hundreds of genes^14, 16^. Rather, our algorithm detects activation as the moment of strong deviation from basal expression levels.

Activation times for *X. tropicalis* ZGA genes in hybrids were similar across replicates (Figure S7A-B), and were correlated with wild-type *X. tropicalis* activation times generated from our algorithm applied to published wild-type data ^17^ (Figure S7C). These results further validate our approach and demonstrate that activation times for *X. tropicalis* genes in hybrids represent a meaningful biological feature of *X. tropicalis* zygotic gene expression timing.

Decreased DNA per cell led to delayed activation of ZGA genes. We found that the mean activation time in cybrids was delayed by ∼21 min (Figure 4B-D), which is near the time of 1 cleavage division cycle in hybrids (∼26 min, Figure 5A). Indeed, the majority of ZGA genes (76.4%; 412/539) exhibited a delay in activation times in cybrids and only a few genes (1.3%; 7/539) advanced their activation times similarly in both replicates (Figure S7E-G). The shift in activation times did not depend upon a gene’s expression level (Figure 4E) and there were no clear subpopulations whose activation timing was not sensitive to DNA content (Figure S7H). Thus, by incorporating information from the entire time-series, we show that gene expression occurs both earlier and at a higher level at a given time point in embryos with more non-template DNA.

**Figure 4.**
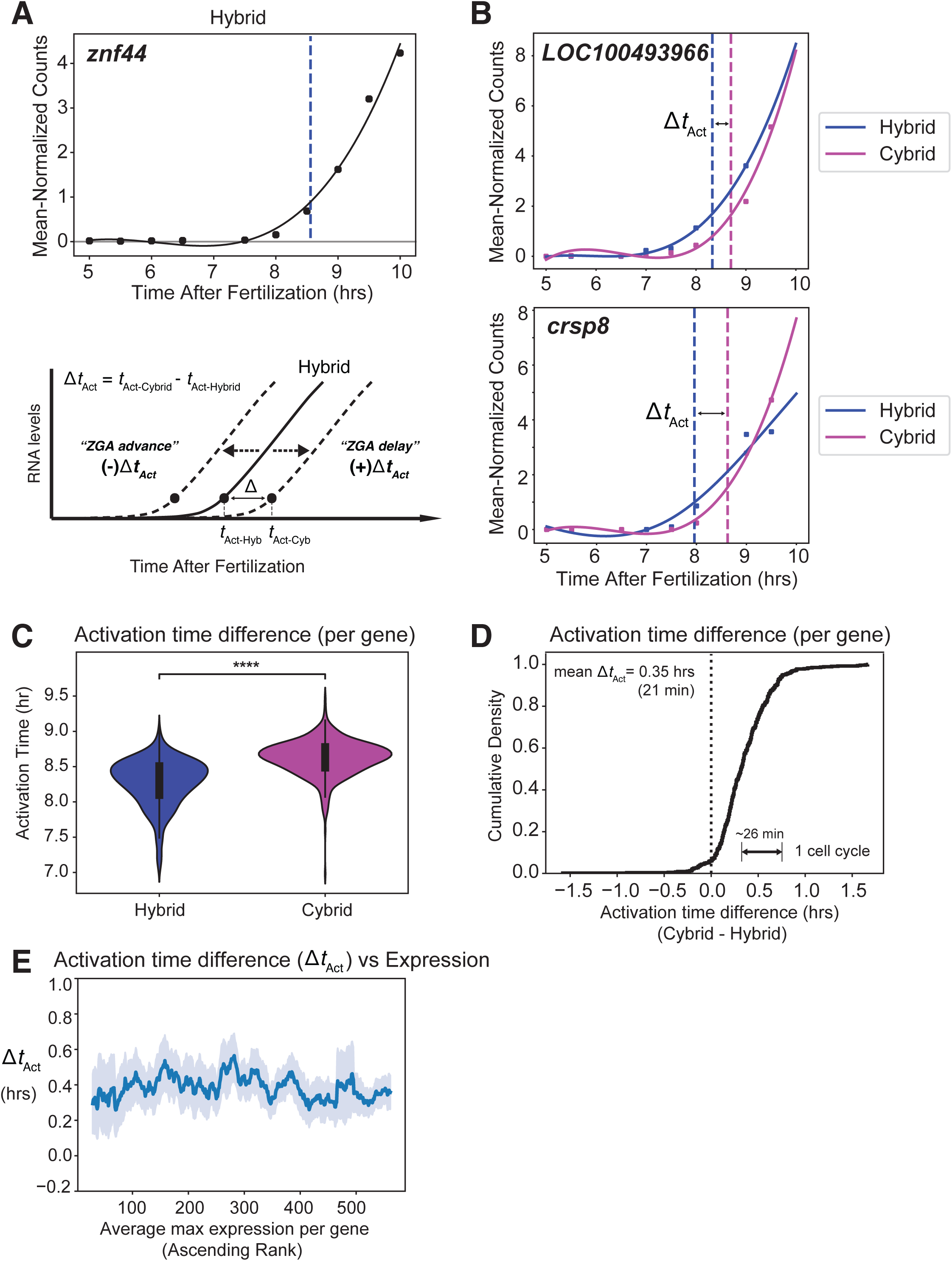
Decreased DNA per cell leads to delayed ZGA gene activation. (A) Smoothing spline fit (black line) of hybrid embryo RNA-seq time-series for example ZGA gene, *znf44*. Black dots denote mean-normalized expression values at each time point. Blue dashed line denotes the estimated “activation point” (see methods and main text). One replicate shown as representative. Schematic below displays activation time approach to compare transcriptomic time-series data. (B) Curve fit and estimated activation times for example ZGA genes, *LOC100493966* and *crsp8*, in hybrid (blue) and cybrid (purple) embryos. The time difference between dashed lines (Cybrid - Hybrid) is the activation time difference, or Δ*t*_Act_. One replicate shown as representative. (C) Violin plot of mean activation time for each gene over two replicates in hybrid and cybrid embryos. Box plot in center shows median, 1^st^ and 3^rd^ quartiles. **** indicates p-value << 10^-10^ (paired sample t-test w/Bonferroni correction). (N=547) (D) Cumulative distribution of the activation time difference (hrs) between cybrids and hybrids. (N=547) (E) Relationship between Δ_Act_ and maximum expression for each gene. Values were binned and smoothed over a 30 min sliding window. Light blue region is standard error. Replicate 3 shown as representative.

**Figure 5.**
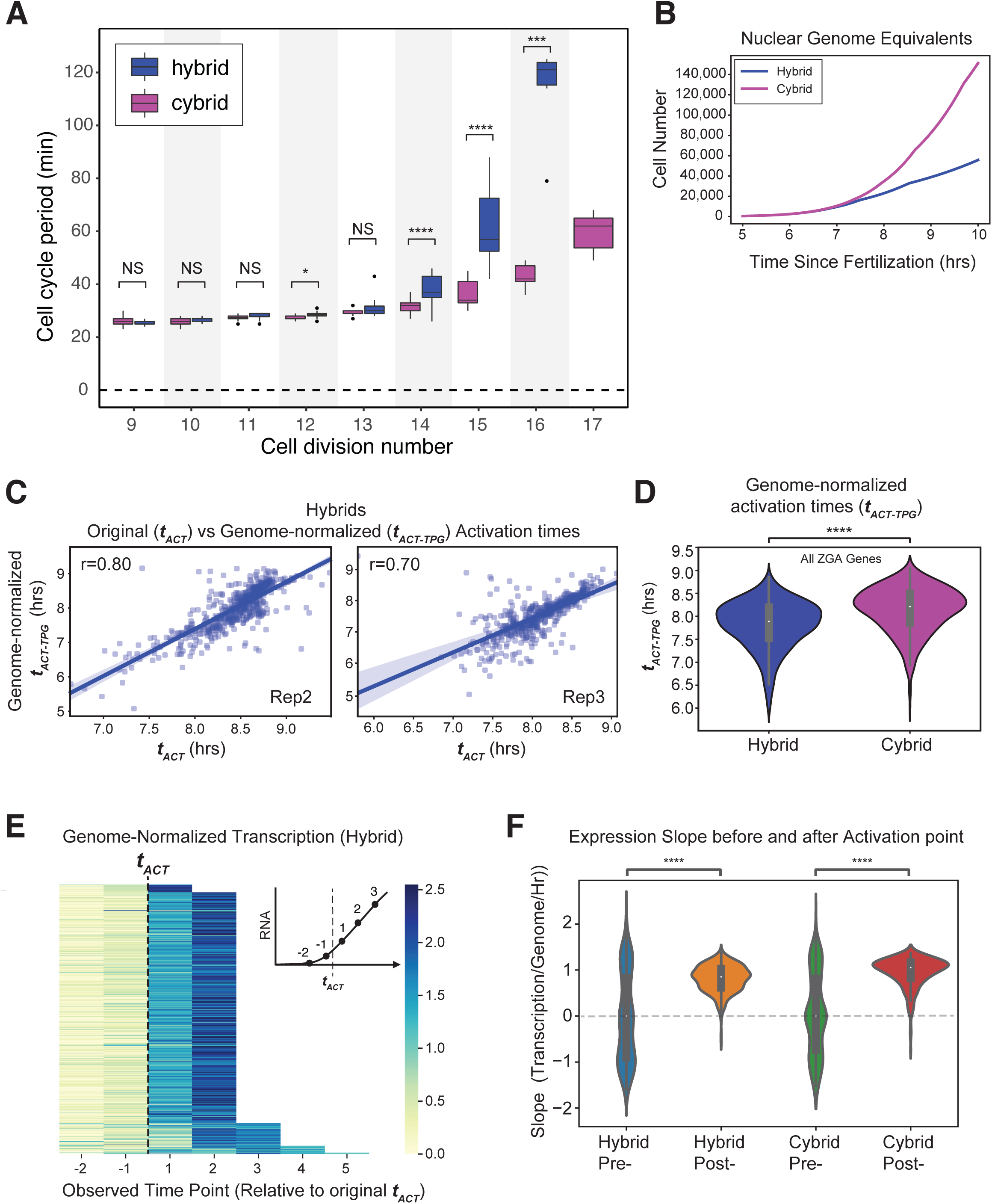
Transcription per genome accelerates through the MBT. (A) Cybrid embryos display early cell cycle lengthening compared with hybrid embryos. Boxplot shows mean inter-cleavage periods for single cells during embryogenesis, in hybrids (blue) and cybrids (red). Values taken from movie (see methods). The box height presents the 25th and 75th percentiles, and the centerline is the median. Whiskers extend to include all non-outlier data points. Single dots are outliers. ns = not significant, or p-value > 0.01 between hybrids and cybrids; * = p-value = 0.0012 for this sample; **** = p-value < 0.0001 between hybrids and cybrids in the same cell cycle (all two-tailed t-test with unequal variance) (B) Line plot showing estimated genome-copy number for hybrids (purple) and cybrids (blue). Data calculated and extended from inter-cleavage division times in panel A. (C) Scatterplot showing relationship between activation time, ***t***_Act_, and genome template number normalized activation time, ***t***_Act-TPG_, for each ZGA gene. Pearson correlation, r=0.80 (replicate 2, left, N=570), r=0.70 (replicate 3, right, N=538). ***t***_Act-TPG_ values were calculated from the genome-normalized time-series using the same smoothing spline fit approach as in Figure 4 (see methods). (D) Comparison of genome-normalized activation times (***t***_Act-TPG_) between hybrids and cybrids. Replicate mean for each condition shown. **** = p-value < 0.0001 (paired sample t-test w/Bonferroni correction). (N=520) (E) Heatmap showing genome-normalized expression values for each gene, aligned at time points adjacent to the original activation time, ***t***_Act_. Legend shows mean-normalized expression values. Inset schematic shows the position of numbered time points relative to ***t***_Act_. (F) Violin plot showing the slope of the genome-template normalized gene expression curves for time points before and after ***t***_Act_. Hybrid values in blue and orange (left); Cybrid values in green and red (right). Note the mean slope changes from ∼0 to ∼1 mean-normalized transcripts per genome per hour, indicating an expression increase above that predicted from genome template number alone.

### DNA content regulates cell cycle duration at the MBT

One hallmark of early development is the transition from rapid and synchronous cell divisions into asynchronous and slower cell cycles. This cell cycle lengthening is regulated by the N/C ratio across many species (reviewed in ^2^) and is likely due to a combination of molecular mechanisms including activation of CHK1, downregulation of CDK1, and titration of replication factors and histones ^37, 39–46^. We assessed whether *Xenopus* embryos with less DNA per cell delay the progressive slowing of the cell cycle by visualizing the animal surface of hybrids and cybrids and measuring multiple single-cell cleavage events across the embryo. Cybrid embryos with 3-fold less DNA have cleavage cell division lengths similar to that of hybrids (26.1 vs 26.0 min, respectively), yet cybrids perform 1-2 more rapid cell division cycles before transitioning to significantly slower division cycles (Figure 5A). Embryos at the 14th and 15th divisions have a mean inter-cleavage period of 32 ± 0.6 min SEM and 36 ± 1.2 min SEM, respectively, in cybrids, compared to 38 ± 1.1 min SEM and 62 ± 3.5 min SEM in hybrids (Figure 5A). Our observations are consistent with N/C ratio control of the cell cycle and the well-established result that haploids with half the DNA of diploids delay cell cycle lengthening by roughly one cycle.

### Transcription per genome accelerates through the MBT

The delayed lengthening of the cell cycle in embryos with less DNA led us to ask whether the increase in transcript levels are due simply to the exponential increase in genome copy number, or whether additional mechanisms contribute to MBT transcript dynamics. We therefore sought to determine how each ZGA gene’s transcription dynamics depend on the number of gene copies per embryo at a given time point. To investigate this, we normalized our transcriptomic time series by the estimated number of cells per embryo at each time point. This was accomplished by fitting a smooth curve to our measurements of cell cycle durations (Figure 5B) (see methods).

Having estimated transcription per gene copy, we sought to determine if our identification of transcriptional activation times was sensitive to the number of gene templates per embryo. We applied our smoothing spline analysis to this genome-normalized transcriptomic profile for each gene, and algorithmically identified activation times (*t*_Act-TPG_) for hybrids and cybrids. Activation times estimated using transcripts-per-genome were in general agreement with those from our analysis without genome normalization (r = 0.80 and r = 0.70 for hybrids, 2 replicates) (Figure 5C). We note that activation times were estimated to be roughly 30 minutes earlier on a gene-by-gene basis in the genome-normalized dataset, suggesting that this approach may more sensitively detect transcriptional activation. Finally, genome-normalized activation times showed a gene expression delay in cybrids with less DNA (Figure 5D). Considering that cybrids will generally have more cells per embryo at a given post-MBT time point (Figure 5B), this result strongly indicates that the observed N/C ratio phenotype is not simply due to cell number differences between hybrids and cybrids.

Our transcription per genome analysis identified an important increase in the transcription rate per gene copy through ZGA. To view this, we centered each ZGA gene’s genome-number normalized transcription time series on the time points before and after the estimated original activation time (*t*_Act_) (Figure 5E). The slope in expression-per-genome-per-hour of ZGA gene population expression profiles is centered around zero before the genome-normalized activation time, with the high variance likely due to much lower expression levels. We then observed a sharp increase in transcription immediately following activation as the slopes significantly increased (Figure 5F). This is consistent with a template number-independent increase in transcript levels across the ZGA. Thus, increased transcription near the MBT is due to an N/C ratio-dependent “boost” to transcription per gene copy in addition to the exponential increase in the genome template after each cell division.

## Discussion

Overall, our results demonstrate that the DNA content in embryos of constant size and maternal environment clearly affects the timing of zygotic transcription. To show this, we used hybrid frog embryos of *X. tropicalis* and *X. laevis* to manipulate embryonic DNA content by 3-fold, while maintaining a constant number of *X. tropicalis* genome templates from which to measure zygotic transcription. The presence of the *tropicalis* genome in the *laevis* maternal environment allowed us to accurately define zygotically activated genes. We compared hybrid embryos with cybrid embryos where the *laevis* genome had been destroyed prior to fertilization using mild UV irradiation so that there is ∼3X less DNA per cell. Broad zygotic genome activation and cell cycle lengthening were delayed in cybrid embryos. Moreover, we did not find evidence of any specific subsets of genes whose zygotic activation did not depend on the N/C-ratio. DNA is therefore a major component of the N/C-ratio that regulates the timing of large-scale zygotic transcription and accelerates gene expression through the MBT in *Xenopus*.

Our experiments inform the debate around the identity of the “N” and “C” components of the N/C-ratio model. The denominator, “C”, appears directly proportional to the cytoplasmic volume. Blastomeres in the same division cycle are either larger or smaller depending on their position in the embryo, and the onset of bulk zygotic transcription occurs earlier in smaller cells and generally depends on cell size rather than the time since fertilization ^30^. The specific aspects of cell size that regulate transcription are unclear, but likely relate to amounts of histones, replication factors, or other molecules whose decreasing cellular amounts can be titrated against the constant amounts of some nuclear factors, N ^4, 37, 39, 41^. Here, our work shows that DNA is a critical component of “N”, while previous work also found a contribution to N from nuclear volume ^28, 29^.

One of the more surprising findings in this work is that we failed to find evidence of a discrete subpopulation of genes whose expression timing at the MBT did not depend on the N/C-ratio. Instead we found that the vast majority of defined ZGA genes delayed their expression in response to 3X less DNA per cell. This result is in contrast to evidence for N/C-ratio- and non-N/C-ratio-regulated gene subsets in *Drosophila* ^26^. Similarly, in another model vertebrate, zebrafish, it was recently reported that at ∼90% of zygotic genes showed some N/C-ratio dependency and ∼10% were N/C-ratio independent ^23^. Taken together, our work indicates that the vast majority of ZGA genes in fast developing vertebrates is regulated by the N/C ratio.

While our work definitively shows that DNA is a crucial component of the global N/C ratio regulating ZGA, the classic N/C-ratio model is incomplete to explain ZGA initiation. For example, the expression of the first gene in zebrafish, miR-430, is unchanged in tetraploid embryos ^47^. In addition, zygotic transcription can be detected prior to the MBT when zebrafish or *Drosophila* embryos are cleavage-arrested at low N/C-ratios ^23, 43^. Moreover, as zygotic genes are activated at distinct times and levels, diverse transcriptional activators must contribute to determining when, where, and how much transcriptional activation takes place for each gene. Thus, many non-mutually exclusive regulatory mechanisms such as basal activator accumulation, Brd4 activity and -mediated histone acetylation of gene promoters, maternal transcription factor translational regulation, and maternal signaling-induced co-factors all likely act in concert to precisely control the broad onset and timing of zygotic genome activation ^14, 19–21, 23, 25, 27, 48, 49^. Here, we conclusively demonstrate that the N/C ratio is also a crucial component regulating ZGA. We propose that the increasing N/C ratio and the exponentially increasing number of genomes globally boosts transcription (Figure 6), but only after chromatin changes and transcription factors have established transcriptional competence or initiated basal transcription in earlier cleavage stages at specific genes.

**Figure 6.**
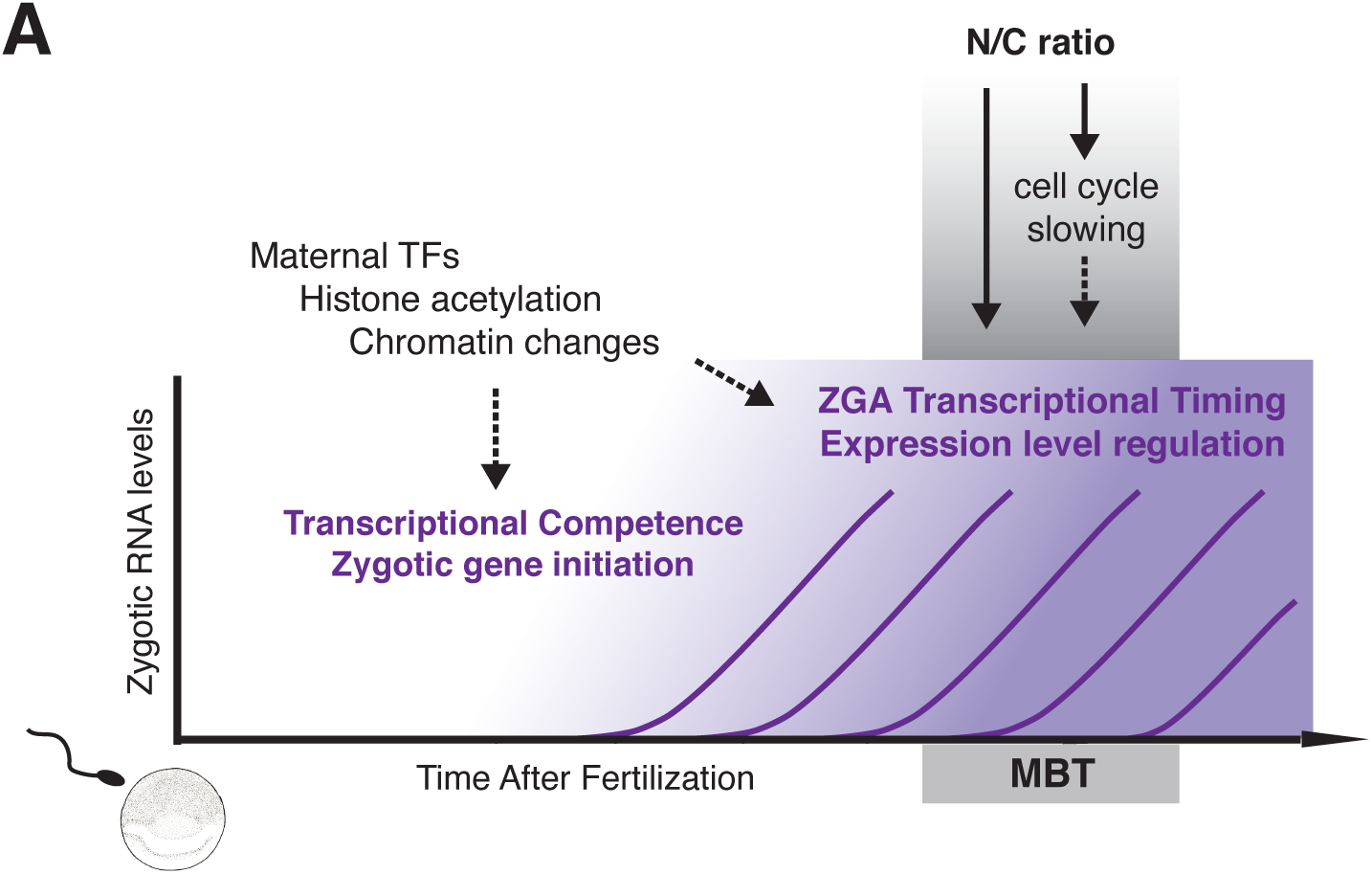
Integrative Model for N/C-ratio control of ZGA timing at the MBT. (A) Schematic of model for ZGA timing. Maternal factors, histone acetylation, and diverse chromatin changes establish transcriptional competence of the embryonic genome, and regulate the transcriptional initiation of up to several hundred genes during cleavage stages prior to the mid-blastula transition ^23, 25^. These standard transcriptional initiation processes are likely required throughout embryo development. A rapid increase in the DNA-to-cytoplasm (N/C) ratio initiates the MBT and regulates ZGA transcriptional timing for the vast majority of zygotic genes at this stage. These non-mutually-exclusive layers of regulation integrate to control the precise initiation and stage-dependent expression level increase of each gene across early embryo development.

## Acknowledgements

We thank Devon Chandler-Brown, Shicong (Mimi) Xie, Charles Limouse, and the Skotheim and Straight labs for critical discussions, and thank Shicong Xie and Amanda Amodeo for comments on the manuscript. We are grateful to Sylvia Choo and Laetitia Parc for technical assistance. We thank Pehr Habury for use of the arc lamp, and Mark Krasnow for use of the stereo-microscope and video camera. The RNA-seq libraries were sequenced at the Stanford Functional Genomics Facility and Novogene. We acknowledge the National Xenopus Resource (RRID:SCR_013731NXR) at the Marine Biological Laboratory (Woods Hole, MA) for supplying inbred Nigerian *X. tropicalis* frogs and training at the *Xenopus* Bioinformatics Workshop. RRK was supported in part by the Stanford Biology Summer Undergraduate Research Program (BSURP). This work was supported by the NIH through grants F32 GM108295 (DJ) and NIH RO1 HD085135 (AFS and JMS).

## Author Contributions

DJ, AFS, and JMS conceived of the research; DJ designed and performed the experiments; DJ and RRK performed computational analysis; DJ and RRK analyzed the data; DJ and JMS wrote the paper with input from all authors. AFS and JMS provided resources and supervised the project.

## Supplemental Figure Legends

**Supplemental Figure 1.**
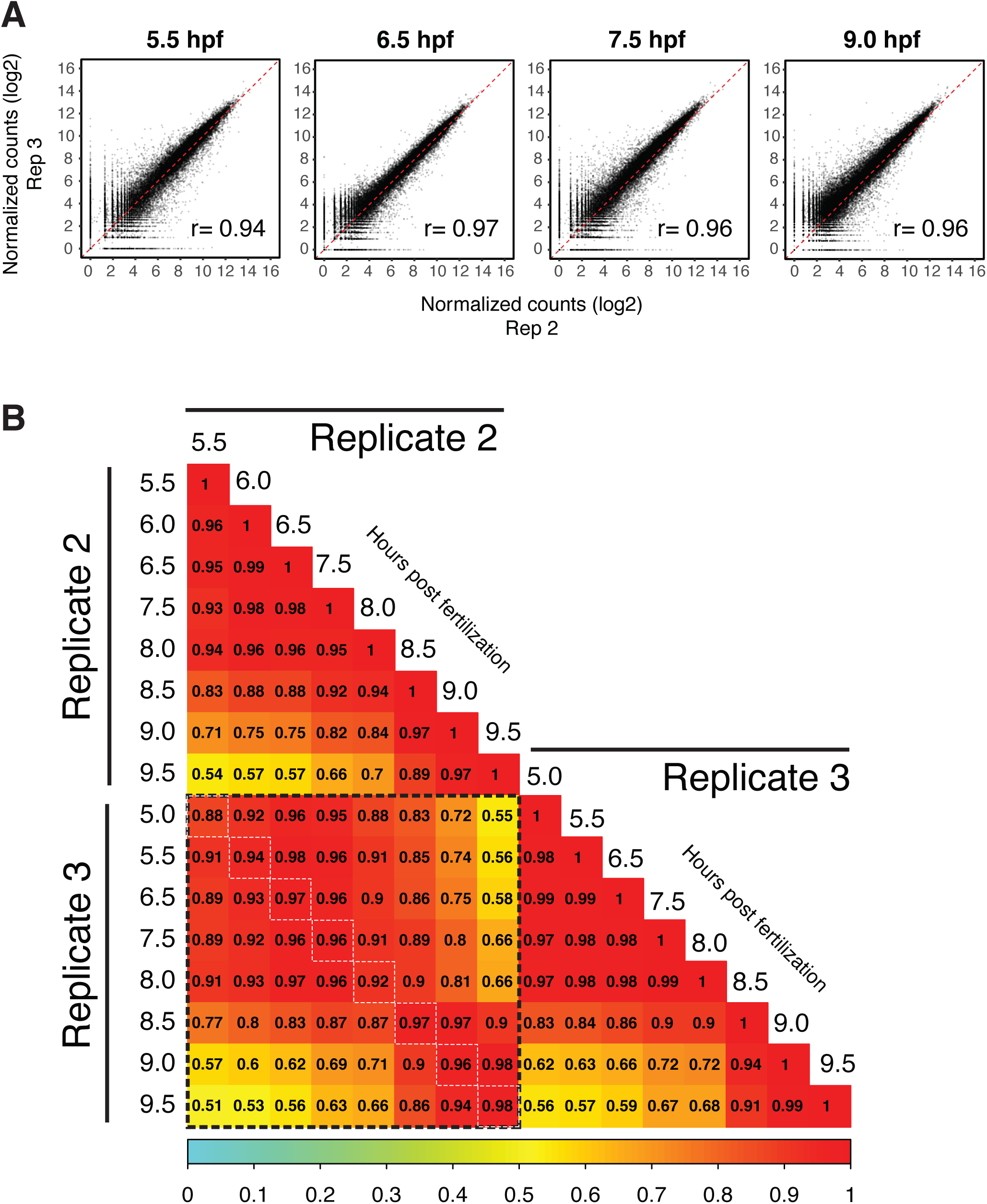
RNA-seq time series and replicate data quality assessment. (A) Scatterplot of gene expression values for all genes in hybrids for 2 replicates indicates high reproducibility. Time points shown include 5.5 hours post fertilization (hpf), 6.5 hpf, 7.5 hpf, and 9.0 hpf. X- and Y-axes show log2(counts) of spike-in normalized RNA-seq gene counts. Dashed diagonal line is X=Y. Pearson correlation shown in inset of each plot. (B) Correlation matrix for replicates 2 and 3 at similar time points in hybrids. Pearson correlation for gene expression values is shown in each box. Color corresponds to Pearson correlation value as indicated in the legend below. To reduce zero-inflated correlation effects, we only considered a gene if it had >40 counts total across the time series in both replicates (mean expression >2.5 counts per time point). White dashed boxes correspond to most highly similar time points between replicates.

**Supplemental Figure 2.**
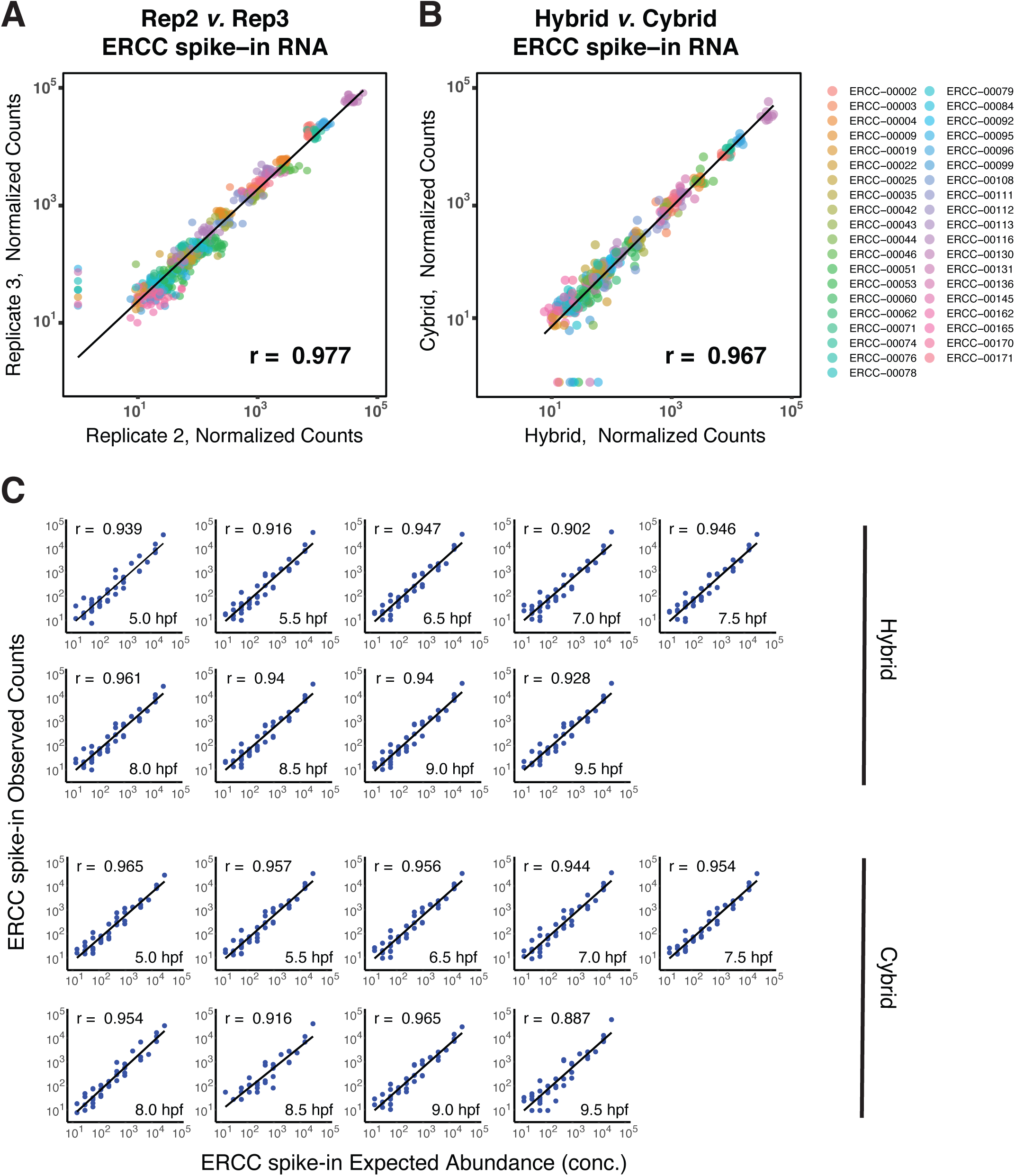
Exogenous spike-in RNA quality assessment. (A) Scatterplot of ERCC (External RNA Controls Consortium) spike-in RNA from two hybrid replicates at all time points shows high reproducibility for each of 38 unique spike-in RNA species used during normalization, as well as the spike-in population. Spike-in RNAs were used to normalize expression counts from all time points and conditions within each replicate (see methods). Each color represents a spike-in RNA species, and each RNA includes the 8 high-quality time points found in both replicates. Black line denotes the linear fit to the data. X- and Y-axes show spike-in normalized RNA-seq counts. ERCC RNAs along the Y-axis (7/38) had low expression in one time point in one replicate that had lower sequencing read depth. Pearson correlation shown in lower right corner (r = 0.977). (B) Scatterplot of ERCC spike-in RNA from the hybrid (high ploidy) and cybrid (low ploidy) conditions reveals consistent expression in both conditions at all time points. Hybrid and Cybrid data are from matched samples from a single egg clutch. Each of 38 unique spike-in RNAs are represented by a single color. Black line denotes the linear fit to the data. Pearson correlation shown in lower right corner (r = 0.97). (C) Scatterplot of observed spike-in RNA counts and their expected abundance based on their known concentrations upon addition to samples. High correlations are present for all time points in both hybrids and cybrids shown for one replicate. Similar correlations were found for the second replicate (data not shown). Each blue dot represents one unique spike-in species. Black line denotes the linear fit to the data. Pearson correlation shown in upper left of each plot.

**Supplemental Figure 3.**
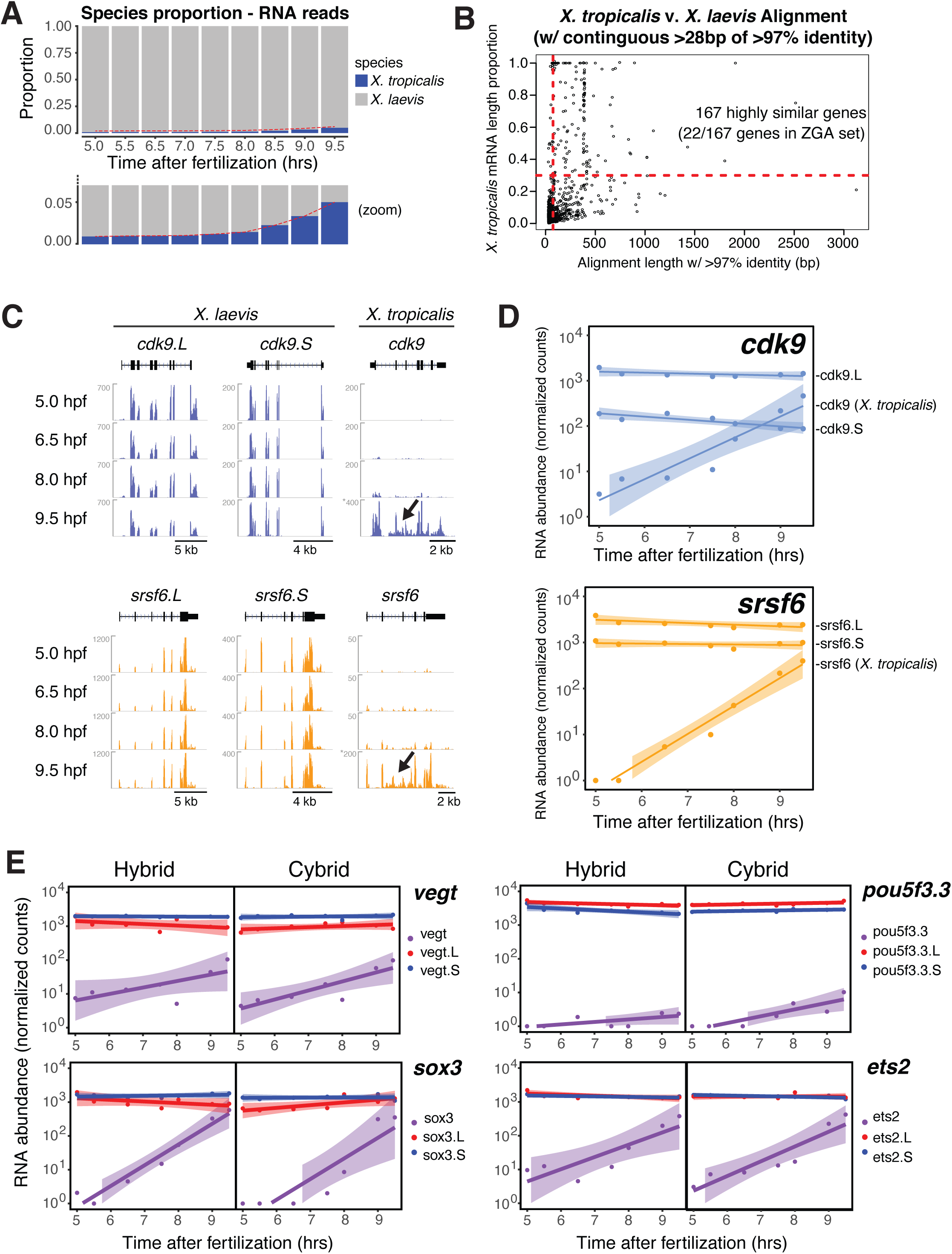
Interspecies alignment quality assessment in hybrids. (A) Bar plot showing proportion of read counts for *X. laevis* and *X. tropicalis* across time points. Color indicates species as in legend. (B) Scatter plot showing results of cross-species BLAST between the *X. laevis* and *X. tropicalis* transcriptomes (mRNA sequences). There were ∼2400 interspecies gene pairs with at least one contiguous nucleotide stretch of >97% identity; each point is one gene pair; genes within a pair are not mutually exclusive, e.g., one *X. laevis* gene could be the top BLAST hit for multiple independent *X. laevis* genes. X-axis shows the length of contiguous nucleotides with >97% identity. Y-axis shows the proportion of each *X. tropicalis* mRNA covered by the contiguous nt region. The blue lines indicate thresholds of 0.3 (Y-axis) and 75 nt (X-axis). All genes (n=22) with at least one stretch of 75 nt with >97% interspecies identity with a length over 30% of the total mRNA were not included in the final ZGA gene set (upper right portion of plot). (C) Genome browser images showing gene expression signal (spike-in normalized read counts) for *X. laevis* and *X. tropicalis* transcripts for *cdk9* and *srsf6*. Signal tracks include merged data from replicates 2 and 3. Read count shown in Y-axis. Signal intensity was originally at base-pair resolution (unbinned) with mean signal over windows shown for clarity. Transcript structure at top in black (boxes = exons; line = introns). Note the lack of intronic signal in maternal *X. laevis* genes that are likely mature, spliced transcripts, and the presence of intronic signal in zygotic *X. tropicalis* genes (black arrow) that likely reflects nascent transcription. *cdk9* (top) and *srsf6* (bottom) shown as representative MZT genes. (D) Expression time-course profiles for *cdk9* and *srsf6*, which have both maternal (*X. laevis*) and zygotic (*X. tropicalis*) mRNA expression in hybrids. The *X. laevis* genome had undergone a duplication and subsequent divergence that results in two subgenomes, short (S) and long (L). The subgenomes include paralogous transcripts for most mRNAs that have high, but not identical, sequence homology. The lack of cross-alignment in the earlier time points indicates that each RNA species (*X. laevis* L, *X. laevis* S, and *X. tropicalis*) can be distinguished. Gene expression was normalized using spike-ins. Points are overlaid with a lowess fit and the associated standard error of the fit. (E) Expression time-course profiles for maternal transcription factors critical for ZGA in *Xenopus* (Gentsch et al., 2019b). *Vegt*, *sox3*, *pou5f3.3*, and *ets2* have both maternal (*X. laevis*) and zygotic (*X. tropicalis*) mRNA expression in hybrids. As in panel C, the lack of cross-alignment in the earlier time points indicates that sequencing reads arising from each species can be distinguished. Gene expression was normalized using spike-ins. Points are overlaid with a lowess fit and the associated standard error of the fit.

**Supplemental Figure 4.**
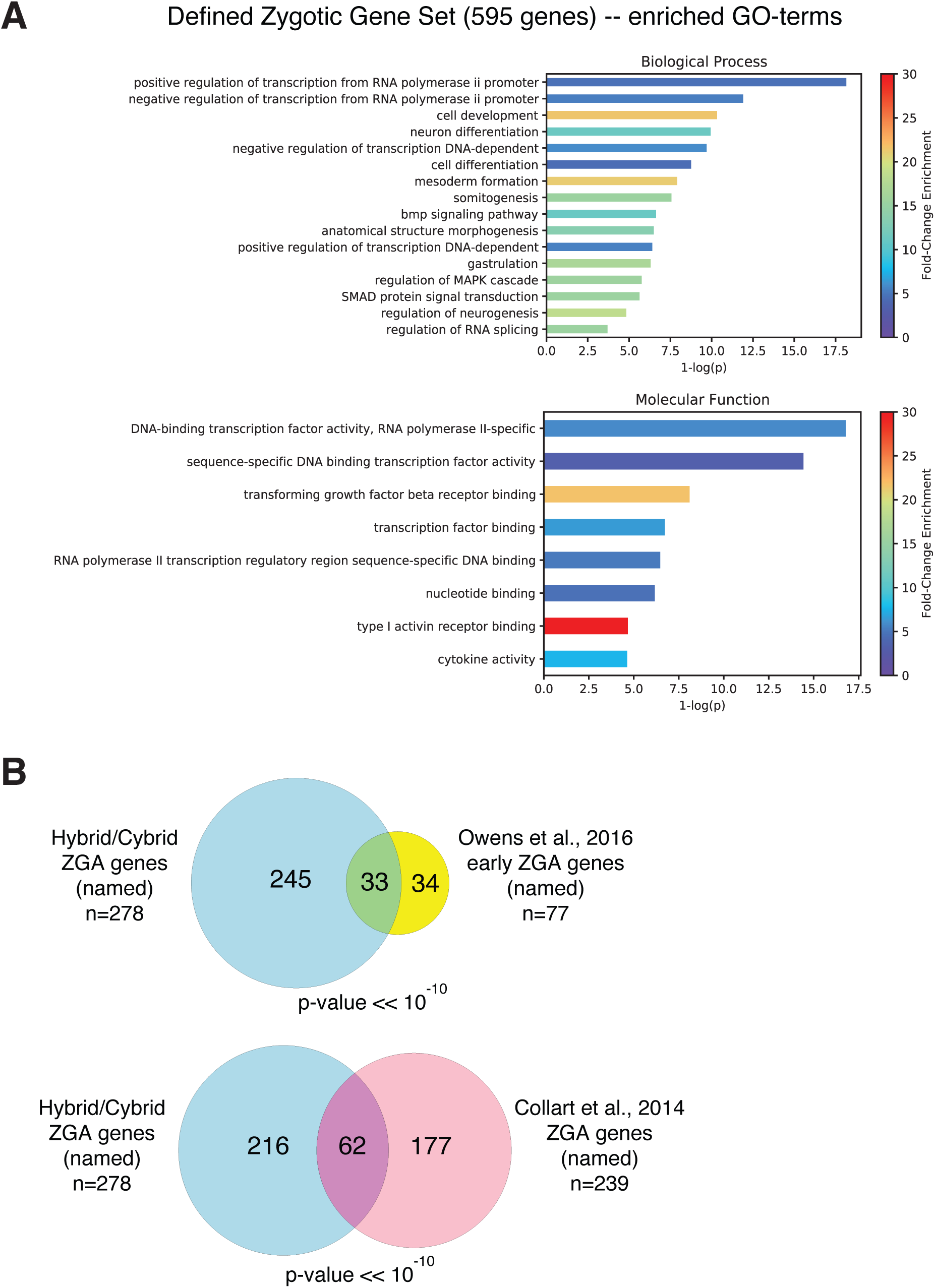
Validation of defined zygotically active genes in hybrids. (A) Gene Ontology (GO) term enrichment of the defined ZGA genes compared to the entire transcriptome. X-axis is −log(p-value), where the p-value was calculated using a hypergeometric test. Fold enrichment of each GO-term in ZGA genes vs. the transcriptome is represented by color as displayed in the legend at right. (B) Venn diagram shows overlap between our hybrid *X. tropicalis* ZGA gene set and wild-type *X. tropicalis* ZGA genes defined in published data sets. Only genes annotated with common names were used to directly compare genes from different genome assemblies and annotations, v7 (Collart et al., 2014; Owens et al., 2016) and v9 (this work).

**Supplemental Figure 5.**
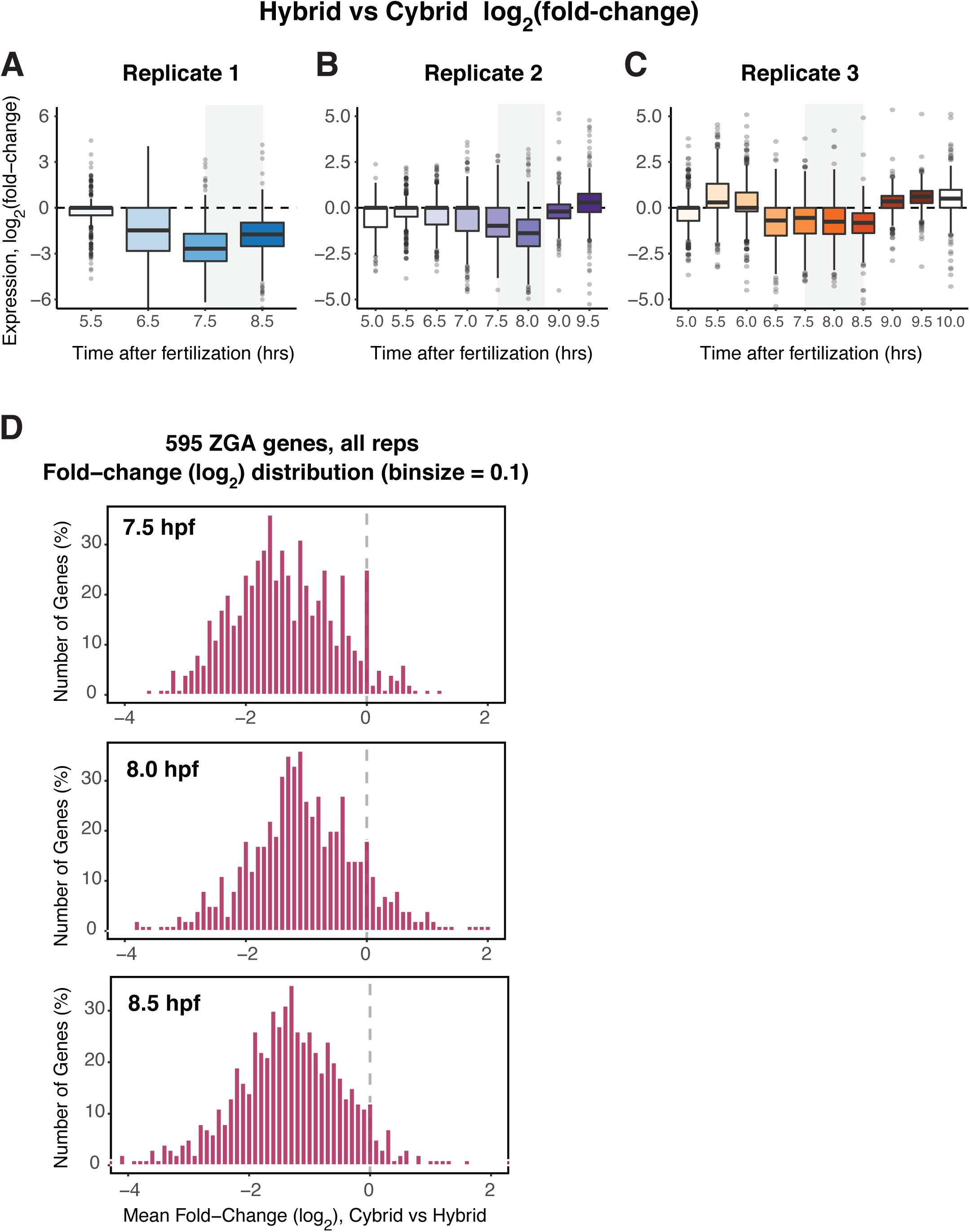
Zygotically activated genes are delayed in embryos with less DNA. (A) Distribution of fold-changes for each ZGA gene’s expression in cybrid relative to hybrid embryos in the replicate 1 time series. Hybrid embryos are the reference, so a decreased fold-change indicates decreased expression in cybrids. For each time point, fold-changes shown are the means of log2-normalized fold-change for all genes from replicate 1. In (A-C), the box shows the mean and 1st and 3rd quartiles. In (A-C) the gray shaded box denotes timing of the MBT. (B) Distribution of fold-changes for each ZGA gene’s expression in the replicate 2 time series. Fold-changes are the means of log2-normalized fold-change for all genes from replicate 2 as in (A). (C) Distribution of fold-changes for each ZGA gene’s expression in the replicate 2 time series. Fold-changes are the means of log2-normalized fold-change for all genes from replicate 3 as in (A). (D) Histograms of fold-changes for defined ZGA genes in all 3 replicates at time points across the MBT (7.5 hpf, top; 8.0 hpf, middle, 8.5 hpf, bottom). Hybrid embryos are the reference, so a decreased fold-change indicates decreased expression in cybrids. For each time point, fold-changes are means of each ZGA gene’s log2-normalized fold-change across 3 replicates (for time points 7.5 hpf and 8.5 hpf) or 2 replicates (for 8.0 hpf).

**Supplemental Figure 6.**
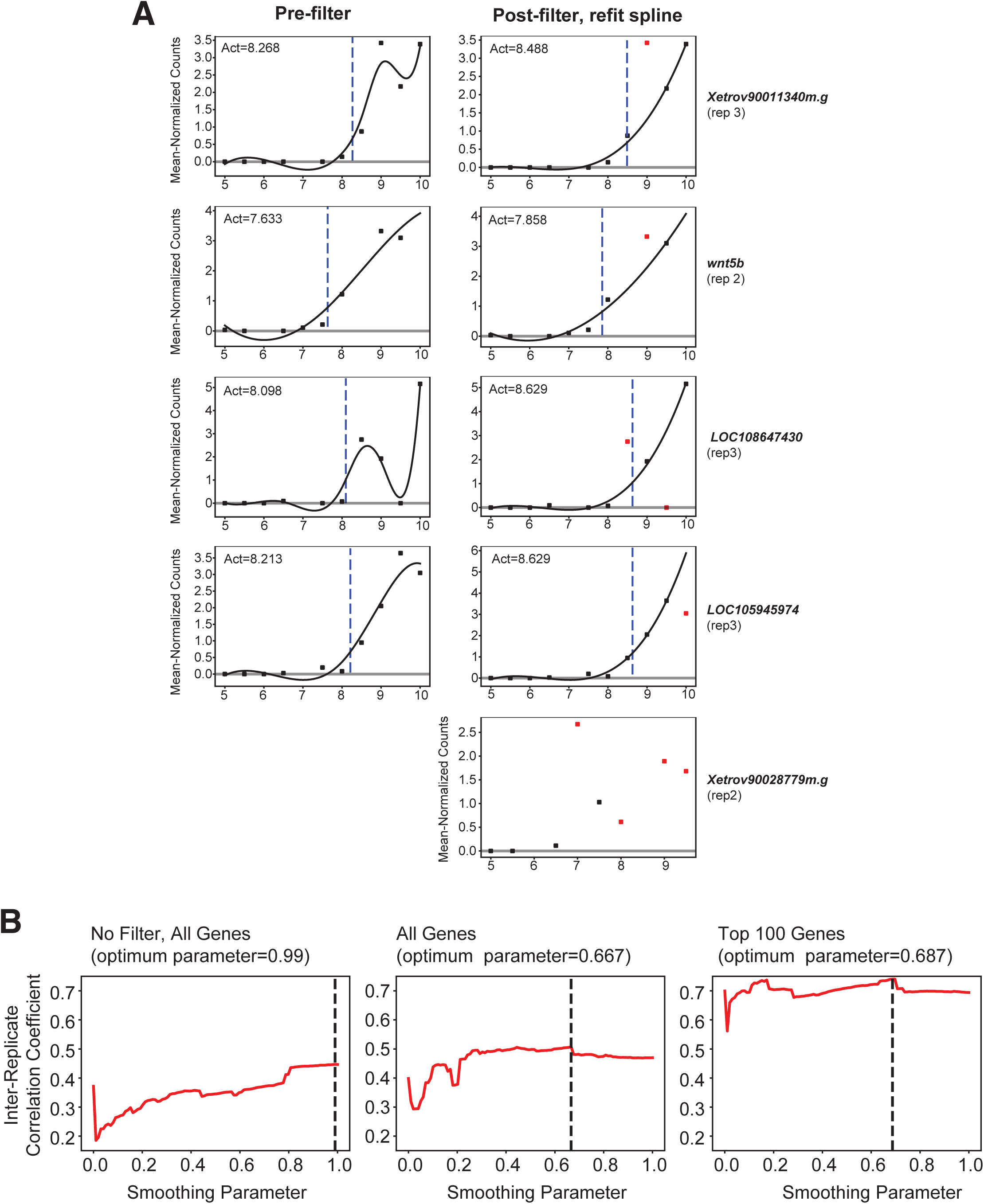
Data processing and smoothing parameter estimation for spline fits of gene expression dynamics. (A) Gene expression profiles showing examples of filtered points. Initial smoothing spline fits were generated using all data points, and any single outlier data point was removed (see methods for precise filter definitions) and a smoothing spline was re-fit to the remaining data (see methods). Black line in each expression profile indicates the re-fit spline. Dashed blue line indicates the inferred “activation time” for each gene. Y-axis expression counts were mean-normalized before applying the filter and spline fit. If 2 or more data points were removed, the gene was not included in downstream analysis. After filtering, 550/595 genes were retained, with activation times determined for 547 genes. (B) Estimation of the optimal smoothing parameter for spline fits. Plots show the correlation coefficient of replicate activation times as a function of the smoothing parameter of the spline fit. Black dashed line indicates the parameter that maximizes inter-replicate correlation. Left plot: Spline fitting with unfiltered data results in low inter-replicate activation time correlation and an inappropriately high and stiff smoothing parameter (i.e., parameters near 1.0 result in a straight-line spline fit across the data set). Middle and Right plots: The optimal parameter is similar whether all genes or the 100 most highly expressed genes are used for the analysis.

**Supplemental Figure 7.**
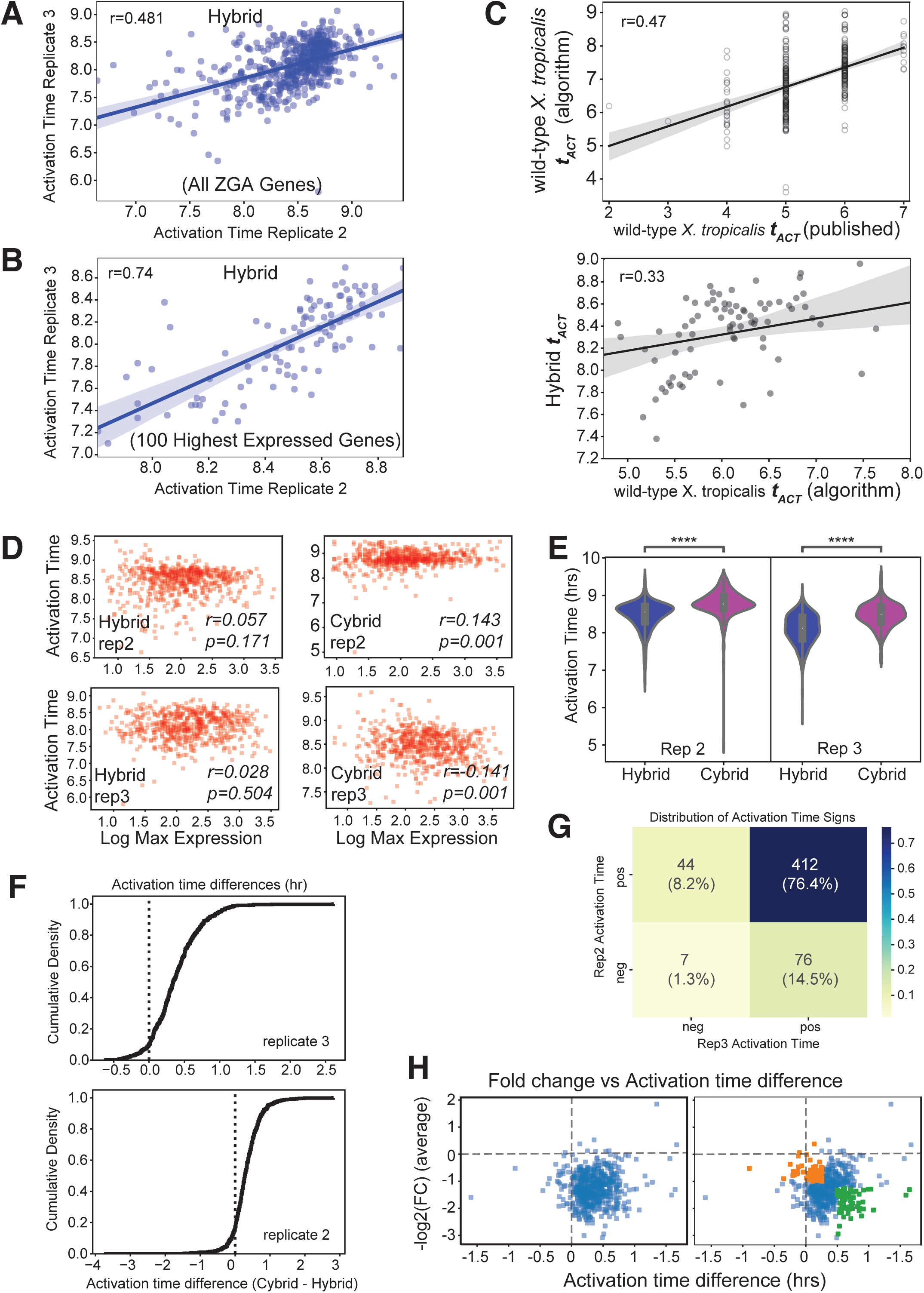
Hybrids and cybrids have different MBT gene activation times for most ZGA genes. (A) Scatterplot of activation times for all ZGA genes comparing replicate 2 and replicate 3. Blue line is linear fit. Pearson correlation in upper left inset. (N=547) (B) Scatterplot of activation times for the 100 most highly expressed (mean expression over time course) ZGA genes comparing replicate 2 and replicate 3. Blue line is linear fit. Pearson correlation in upper left inset. (N=100) (C) Top: Scatterplot of published activation times from wild-type *X. tropicalis* data from an independent time series (Collart et al., 2014) (X-axis) vs activation times from our algorithm applied to this same data. Named genes common to both data sets were included. Bottom: Plot compares our hybrid activation times to algorithmically determined wild-type *X. tropicalis* activation times from (Collart et al., 2014). (N=82) (D) Scatterplot compares activation time and expression level (log2 of the maximum expression across the time course) for each ZGA gene in both hybrid and cybrid conditions in replicates 2 and 3. Pearson correlation and p-value shown in lower right inset of each plot. (rep2, N=575; rep3, N=562) (E) Violin plot of mean activation time for each gene in each replicate in hybrid and cybrid embryos. Same original data as in Figure 4C, but separated by replicates. Box plot in center shows mean, 1st and 3rd quartiles. **** indicates p-value < 10^-10^ (paired sample t-test w/Bonferroni correction). (rep2, N=575; rep3, N=562) (F) Cumulative distribution of the activation time difference (hrs), Δ*t*_Act_, between cybrids and hybrids for each replicate. Same original data as in Figure 4D, but separated by replicates. Dashed black line indicates X=0 (no difference). (rep2, N=575; rep3, N=562) (G) Display of the distribution of the sign of Δ*t*_Act_ across both replicates. A positive sign indicates that cybrid gene expression was less than that of hybrids (i.e., delayed). Percent of ZGA genes indicated by color as shown in legend. (N=539) (H) Scatter plot of mean Δ*t*_Act_ and mean log2(FC) for the 7.5, 8.0, and 8.5 hpf time points for each ZGA gene, all replicates. Negative fold-change and positive Δ*t*_Act_ indicate less expression in cybrids vs hybrids and delayed gene expression activation. Right panel: genes colored are above (green) or below (orange) joint thresholds for Δ*t*_Act_ and mean log2(FC).

## RESOURCE AVAILABILITY

### Lead Contact

Further information and requests for resources and reagents should be directed to and will be fulfilled by the Lead Contact, Jan Skotheim.

### Materials Availability

This study did not generate new unique reagents.

### Data and Code availability

Sequencing read data generated during this study are available at NCBI Gene Expression Omnibus, GSEXXXXX, [weblink].

## KEY RESOURCES TABLE

| REAGENT or RESOURCE | SOURCE | IDENTIFIER |
| --- | --- | --- |
| **Antibodies** |  |  |
| Rabbit anti-xCENP-A | Aaron Straight, Stanford University | Milks et al., 2009 |
| AlexaFluor 568-Donkey anti-Rabbit IgG | ThermoFisher Scientific | Cat# A10042 |
| **Biological Samples** |  |  |
| *Xenopus laevis* / *Xenopus tropicalis* hybrid embryos | This paper |  |
| *Xenopus laevis* embryos | This paper |  |
| **Chemicals, Peptides, and Recombinant Proteins** |  |  |
| Tricaine methanesulfonate (MS-222/Tricaine-S) | Syndel (Mfr.) | Cat# NC0342409 (Fisher Scientific) |
| 8-Methoxypsoralen | Millipore Sigma | M3501-1G |
| TriPure Isolation Reagent (Roche, Mfr.) | Millipore Sigma | Cat# 11667165001 |
| TURBO DNase | ThermoFisher | Cat# AM2238 |
| ERCC RNA Spike-In Mix (Mix 1) | ThermoFisher | Cat# 4456653 |
| L-cysteine free base | MP Biomedicals | Cat# 101444 |
| Pregnant Mare Serum Gonadotropin (PMSG) |  |  |
| Human Chorionic Gonadotropin (hCG), CHORULON | Merck Animal Health / Intervet | CH-475-1 |
| **Critical Commercial Assays** |  |  |
| Script-seq v2 RNA-seq Library Preparation Kit | Illumina (epicentre) | Cat# SSV21124 |
| Script-seq Ribo-Zero™ rRNA Removal Kit | Illumina (epicentre) | Cat# SCL24H |
| Script-seq Index Primers, Set 1 and Set 2 | Illumina (epicentre) | Cat# RSBC10948<br>Cat# SSIP1202 |
| SuperScript III First-Strand Synthesis System for RT-PCR | ThermoFisher | Cat# 18080-051 |
| SYBR GreenER<br>qPCR SuperMix<br>for ABI PRISM | ThermoFisher | Cat# 11760-500 |
| RNAClean XP<br>SPRI beads | Beckman Coulter | Cat# A63987 |
| RNeasy<br>MinElute<br>Cleanup Kit | Qiagen | Cat# 74204 |
| RNA Clean &<br>Concentrator -<br>25 | Zymo Research | Cat# R1018 |
| RNA Clean &<br>Concentrator - 5 | Zymo Research | Cat# R1013 |
| **Equipment** |  |  |
| Hg Arc lamp and<br>500 Watt power<br>supply | Oriel Instruments |  |
| **Deposited Data** |  |  |
| RNA-seq of<br>hybrid and cybrid<br>embryo RNA | This paper | GEO: GSEXXXX |
| **Experimental Models: Organisms/Strains** |  |  |
| *Xenopus laevis*:<br>adult female | Nasco | Cat# LM00535 |
| *Xenopus laevis*:<br>adult male | Nasco | Cat # LM00715 |
| *Xenopus*<br>*tropicalis*: adult<br>female,<br>*Xtr.NigerianSt5*<br>*49* | National Xenopus<br>Resource (NXR) | NXR_1018, female |
| *Xenopus tropicalis*: adult male,<br>*Xtr.NigerianSt549* | NXR | NXR_1018, male |
| **Software and Algorithms** |  |  |
| FastQC v0.11.8 |  | https://www.bioinformatics.babraham.ac.uk/projects/fastqc/ |
| Trimmomatic v0.38 | 50 | http://www.usadellab.org/cms/?page=trimmomatic |
| bowtie2 v2.3.4.1 | 51 | http://bowtie-bio.sourceforge.net/bowtie2/index.shtml |
| Kallisto v0.44.0 | 52 | https://pachterlab.github.io/kallisto |
| STAR v2.5.4b | 53 | https://github.com/alexdobin/STAR |
| featureCounts v1.6.3 | 54 | http://subread.sourceforge.net/ |
| Samtools v1.8 | 55 | http://www.htslib.org/ |
| Bedtools v2.27.1 | 56 | https://bedtools.readthedocs.io/en/latest/<br>https://github.com/arq5x/bedtools2/releases |
| python | https://www.python.org/ | https://www.python.org/ |
| Scipy v1.5.2 | https://docs.scipy.org/ | https://docs.scipy.org/ |
| Rstudio | www.rstudio.com | www.rstudio.com |
| BLAST | 57 | https://blast.ncbi.nlm.nih.gov/Blast.cgi |
| Integrated Genomics Viewer (IGV) (v2.8.13) | Robinson et al., 2011 | http://software.broadinstitute.org/software/igv/ |
| ImageJ |  | https://imagej.nih.gov/ij/ |
| Adobe Illustrator CC 2020 |  | www.adobe.com |
| ImagePro |  | https://www.mediacy.com/imagepro |

## EXPERIMENTAL MODEL AND SUBJECT DETAILS

Adult, sexually mature wild-type *Xenopus laevis* females (1-3 years old) were obtained from Nasco (LM00535MX). Adult sexually mature male and female inbred J-strain *Xenopus laevis* were obtained from the National *Xenopus* Resource (NXR, Woods Hole, MA; NXR_0024). Adult, sexually mature inbred Nigerian strain *X. tropicalis* males (7 months to 2 years old) were obtained from NXR (NXR_1018). All frogs were housed and maintained in the Stanford Aquatic Facility staffed by the Veterinary Service Center. *X. laevis* were housed at 18°C with a 12/12 hour light/dark cycle, and frogs were fed twice weekly *X. tropicalis* were housed at 26-28°C. Animal work was carried out in accordance with the guidelines of the Stanford University Administrative Panel on Laboratory Animal Care (APLAC).

## METHOD DETAILS

### Fertilization of *X. laevis* females with *X. laevis* sperm

For ovulation, female *X. laevis* were primed 2-14 days before ovulation by subcutaneous injection of 50 U pregnant mare serum gonadotropin (PMSG; Sigma) at the dorsal lymph sac, with ovulation induced 12-14 hours before egg collection with injection of 500 U human chorionic gonadotropin (hCG; Chorulon). *X. laevis* had a minimum resting period of 5 months between ovulations. During ovulation, frogs were housed individually in 2 L 1X Marc’s modified Ringer’s buffer (MMR; 6 mM Na-HEPES, pH 7.8, 0.1 mM EDTA, 100 mM NaCl, 2 mM KCl, 1 mM MgCl_2_, and 2 mM CaCl_2_) at 17°C. Eggs for *in vitro* fertilization (IVF) were collected by gentle massaging of the female *X. laevis* abdomen into a drop of 1X MMR into a XX cm round glass dish, with excess liquid removed. To obtain *X. laevis* testes, males were euthanized in 2g/L Tricaine-S (MS222) and 5 mM sodium bicarbonate in frog tank water, and testes were immediately dissected, cleaned, and placed into High Salt MBS (20 mM NaCl, 0.7 mM CaCl_2_, 1X Modified Barth’s Saline-MBS) at 4°C and were then either used fresh or were stored at 4°C for up to 7 days. Fertilization was performed by massicating ∼⅓ of one testes in a 1.5 µl eppendorf tube in 200 µl High Salt MBS and pipetting the liquid over the eggs. The remaining testis was used to gently move the eggs into a monolayer. After 3 minutes, excess testes pieces were removed and the dish flooded with 0.1X MMR, and this time was noted as the fertilization time (t=0 hours post fertilization; hpf).

### Fertilization of hybrid *X. laevis* eggs with *X. tropicalis* sperm

Ovulation and egg collection of female *X. laevis* for hybrid generation was performed as above, but with eggs collected in 10 cm plastic petri dishes. To obtain *X. tropicalis* testes, males were euthanized in frog tank water and 2g/L Tricaine-S (MS222) and 5 mM sodium bicarbonate prior to dissection. Testes were obtained immediately through dissection, cleaned and placed into 1X MMR at room temperature for fresh use in fertilization. Eggs used for hybrids were set aside for 15-20 minutes while the eggs used for cybrids were manually selected (see below). Both testes from a single male were masticated in a 1.5 µl Eppendorf tube with 400µl 1X MMR and added to the *X. laevis* eggs. The petri dish was tilted ∼15% and excess testes pieces were rubbed over the eggs. After 3 minutes, the dish was flooded with ddH20 and this time was noted as the fertilization time (t=0 hpf). After 5 minutes, the ddH20 was gently exchanged for 0.1X MMR. The jelly coat was removed 25 minutes after fertilization by incubating eggs in 1X MMR + 2% (w/v) L-cysteine, pH = 8.0 for 2-4 minutes. Embryos were then washed 3-4 times with 0.33X MMR and incubated at 0.33X MMR at room temperature (21°C) until sample collection. Embryos in all conditions were monitored and embryos with aberrant cleavages or any signs of poor health were removed.

### UV-irradiation to generate Cybrids

To generate cybrid embryos that lack maternal DNA, a subset of the *X. laevis* eggs collected for hybrids were selected for UV-irradiation prior to fertilization. Embryos to be irradiated were treated with 25 µM of 8-methoxypsoralen in 1X MMR for 10-15 minutes before UV-irradiation, which has been shown to aid in complete *X. laevis* genome cross-linking without adversely affecting the health of the embryo or ability to develop into the tadpole stage ^36^. Prior to making cybrids, we tested different UV wavelengths, crosslinking conditions, and UV-light durations on *X. laevis* eggs to identify the optimum conditions for which maternal egg DNA is completely absent from resulting embryos, but where the eggs remain healthy, have a fertilization efficiency similar to that of unirradiated eggs using both *X. laevis* and *X. tropicalis* testes, and match the survival rates to tadpoles in *X. laevis* x *X. laevis* crosses generated using irradiated *X. laevis* sperm instead of the egg (data not shown). For cybrid fertilizations, 50-60 eggs were individually transferred to a dry 6 cm plastic petri dish using a Dumont #55 forceps to gently hold the jelly coat. Individual eggs were surrounded by a small drop of 0.3X MMR, and manually adjusted with the pigmented side up (*i.e.*, germinal vesicle near the top), with care taken to touch only the jelly coat and avoid direct contact with the egg. The petri dish was filled with 0.3X MMR. Eggs were irradiated by placing the 6 cm petri dish in a dark chamber for 2 minutes under a 500 Watt Hg arc lamp (Oriel Instruments) producing a downward facing focused beam with diameter of ∼8 cm such that every egg in the dish was covered by the UV light. The UV light was filtered to only allow 300-400 nm wavelength light onto the sample with a peak wavelength of ∼350 nm. Following irradiation, the petri dish solution was immediately exchanged for 1X MMR. The small petri dish was then placed in a large glass dish. Fertilization was achieved as with hybrids, with both hybrid and cybrid eggs from the same clutch receiving an equal portion (∼200 µl) of the testes/MMR solution from the same male. Fertilizations of hybrid and cybrids were synchronized and embryos de-jellied as described above.

### Embryo time course sample collection for RNA-seq

Hybrid and Cybrid embryos from the same fertilization event were collected in stage-matched samples. 3 replicates were obtained from independent females on different days: replicate 1 consisted of 4 time points spaced 1 hour apart; replicates 2 and 3 were 9-10 time points sampled every 30 min. Embryos were assessed at all stages after fertilization and any embryos with even minor aberrant cleavages were removed (always <10% of embryos in each condition). At each time point, 5 embryos were gently transferred to a 1.5 ml Eppendorf tube using a plastic Pasteur pipet cut with a wide bore tip such that embryos never touched the pipet tip. Embryos were allowed to settle, and excess MMR removed. Embryos were washed quickly with ∼1.5 ml cold RNase-free embryo wash buffer (EWB; 10 mM K-HEPES, pH 7.7, 100mM KCl, 1 mM MgCl_2_, 0.1 mM CaCl_2_, 50mM sucrose). Immediately after removal of EWB, 500 µl of Tripure was added to each tube, which was then vortexed for 30 seconds to homogenize the embryo until no visible pieces remained. 400 µl additional Tripure was added and samples were flash frozen in liquid nitrogen and stored at −80°C until RNA extraction.

### Karyotyping of hybrids and cybrids by Immunohistochemistry of Centromeres

Embryos were karyotyped using protocols from the Grainger lab (University of Virginia) and ^58^ with modifications. Hybrid embryos were used at 40-66 hours post fertilization (NF stage 32-40), and Cybrid embryos that had stopped development after neurulation were taken at 24-36 hours with all non-viable tissue removed. For both, embryos were anesthetized (2g / L MS-222 and 2g / L Sodium Bicarbonate) and rinsed in 0.33X MMR. The yolky ventral portion was removed, and embryos were transferred to ddH2O and allowed to stand for 20 min. Dorsal halves were pipetted into an Eppendorf tube with 60% acetic acid in ddH2O for 5 min, then placed on a positively charged slide with excess liquid removed and covered with a large coverslip. Slides were covered with a paper towel, pressed using a ∼20kg lead brick for 5 minutes, and then placed on dry ice for 5 minutes. The coverslip was removed, and samples fixed in 3.7% paraformaldehyde (formalin) in dilution buffer (80 mM K-PIPES pH 6.8, 1 mM MgCl2, 1 mM EGTA, 30% glycerol, 150 mM KCl, 0.5% Triton-X 100) for 15 minutes in a humid chamber. After 3 rinses, the sample was blocked for 30 minutes in antibody dilution buffer (AbDil; 20 mM Tris-HCl, pH 7.4, 150 mM NaCl with 0.1% Triton X-100, and 2% bovine serum albumin). Samples were rinsed twice and exposed to 10 µg/mL Hoechst 33258 and 1μg/mL rabbit-anti-xCENP-A diluted in AbDil for 1 hour, washed three times with AbDil, and then exposed to 1 μg/mL AlexaFluor 568-conjugated goat anti-rabbit secondary antibodies (Life Technologies) for 1 hour. Slides were rinsed 3 times, mounted in 100 µl 70% Glycerol in 1X PBS, sealed under a coverslip with clear nail polish, and stored at −4°C until imaging.

### Centromere Imaging and Quantification

Imaging was performed on an IX70 Olympus microscope with a DeltaVision system (Applied Precision) a Sedat quad-pass filter set (Semrock) and monochromatic solid-state illuminators, controlled via softWoRx 4.1.0 software (Applied Precision). Images of nuclei from neurulated embryos were acquired using a 60x 1.4 NA Plan Apochromat oil immersion lens (Olympus), and a charge-coupled device camera (CoolSNAP HQ; Photometrics) and digitized to 16 bits. For each nucleus, 10-30 Z-sections at 0.2 μm intervals were taken. Centromeres were counted manually in maximum intensity projections. Ambiguous centromeres were counted by looking through the original stack in imageJ. Displayed images of nuclei are maximum intensity projections (Figure 1).

### Imaging and quantification of cell cycles

To measure cell cycle duration during embryogenesis, we generated and analyzed movies as described in ^37^. Briefly, developing embryos were placed in 0.33X MMR. Time-lapse images were captured every 1 minute on a Leica MZ16FA Stereomicroscope at 24X magnification using Image-Pro software. Movies began at the 8- or 16-cell stage, and divisions were counted to determine the frame number of the eighth cleavage. Then, ∼30 individual cells per embryo were selected from the visible portion (primarily animal side) of each embryo after the eighth cleavage. Inter-cleavage periods were determined by manually tracking single cells and noting the frame number at which the cleavage furrow visibly transected the entire cell. When daughter cells did not divide concurrently, the division time of the earliest dividing daughter was used, and that cell was followed for the remainder of the movie. In cells where the complete cleavage could not be observed, *e.g.*, in cases where the cleavage plane did not intersect with the embryo surface, the cell was omitted from analysis.

### Embryo RNA processing for RNA-seq

RNase-free reagents were used at all steps after embryo collection. Embryo samples in Tripure reagent were thawed at room temperature and placed on ice. ERCC spike-in RNAs (4456653; ThermoFisher Scientific) were added at a 1:40 dilution of stock at 1µl per embryo (5 embryos and 5 µl diluted ERCC RNA per sample) to the samples in Tripure and mixed gently by inversion followed by 2 seconds of vortexing. RNA was extracted by adding 200 µl chloroform to the thawed embryos in 1 ml of Tripure, incubating for 10 minutes at room temperature, and centrifuging at 12,000 x g for 15 minutes at 4°C. The aqueous phase was transferred to a fresh tube with 500 µl of 4°C isopropanol, mixed gently by inversion, and centrifuged at 12,000 x g for 10 minutes at 4°C. The resulting pellet was washed with 1 ml 75% EtOH, centrifuged for 7,500 x g for 5 minutes at 4°C, and the RNA pellet was air dried and resuspended in 20 μl DEPC-treated H20. Following extraction, RNA concentrations were measured by nanodrop and total RNA integrity assayed using an Agilent Bioanalyzer. All RNAs had a RIN (RNA integrity number) greater than 9.0. RNA (10 μg) was then treated with DNase (TURBO DNase; Ambion) for 1 hour at room temperature followed by isolation with a minElute RNA Cleanup Kit (Qiagen). 5 μg DNase-treated RNA was used as input for ribosomal depletion using the Epicentre (Illumina) Ribo-Zero ribosomal depletion kit (Cat# SCL24H). Ribosomal RNAs were depleted and RNA quality was assessed post-depletion using a bioanalyzer to ensure loss of 18S and 28S rRNA. rRNA-depleted RNA was then purified using Ampure SPRI beads with a 1.8X bead to sample ratio.

### RNA-seq library preparation and sequencing

RNA was converted to cDNA libraries ready for sequencing using the stand-specific Script-seq V2 RNA-seq kit (Illumina). cDNAs were generated, amplified, and indexed according to the manufacturer’s instructions. 14 PCR cycles were used to amplify libraries and samples were indexed for multiplexing. Final library concentrations were determined by qPCR using custom primers and exogenously added PhiX DNA (Illumina) to generate a concentration reference curve. To reduce lane effects on individual samples within a replicate, indexed libraries from each time point and embryo treatment condition (hybrid and cybrid) were pooled into a single replicate sample. Each of the 3 biological replicate libraries was sequenced at low read depth on a MiSeq (2 x PE75) at the Stanford Functional Genomics Facility to assess quality, and later on 6 lanes of the HiSeq4000 (2 x PE75 and 2 x PE150) at NovoGene (Sacramento, CA). All reported analysis was generated using the HiSeq dataset.

### RNA-seq read processing

Paired-end RNA-seq reads were assessed for quality using fastQC. Reads from independent sequencing runs for each sample were merged, and reads from 150 nt paired-end runs were trimmed to 75 nt. Adapters were trimmed and low quality sequences removed using trimmomatic and the requisite Illumina adapter sequences (trimmomatic v0.38; PE -threads 2 ILLUMINACLIP:Illumina_TrueSeq_Adapters_PE.fa:2:30:10 HEADCROP:1 MINLEN:31). Adapter-trimmed reads were filtered for any remaining residual rRNA, mitochondrial RNA, and tRNA by aligning to a curated FASTA file containing these sequences, using bowtie2 (v; parameters: --local -X 2000 -p8 --fr --norc -t -x --dovetail --al-conc-gz --un-conc-gz) (Langmead and Salzberg, 2012). Mitochondrial and rRNA sequences were obtained from NCBI and Xenbase and tRNA sequences from the *X. tropicalis* v9.1 transcriptome (XENTR_9.1.transcripts.fa). Reads aligning to the *X. tropicalis ccdc50* gene were also filtered due to homology to highly abundant sequence(s) and extreme read count values up to an order of magnitude higher than that of the next-highest expressed mRNA. All other mRNA genes in both frog species had expression counts within the expected range. Unaligned paired reads depleted for the above RNAs were retained and used as input for hybrid mRNA transcriptome alignment described below.

### Sequencing read alignment to transcriptomes

To distinguish *X. laevis* and *X. tropicalis* transcripts, we first merged both species’ transcriptomes into a composite hybrid transcriptome, consisting of the *X. laevis* v9.1 (XL_9.1_v1.8.3.2.primaryTranscripts.fa) and *X. tropicalis* v9.1 (XENTR_9.1.transcripts.fa), transcriptomes obtained from the Xenbase FTP site (http://ftp.xenbase.org/pub/Genomics/JGI/), and the ERCC spike-in RNA FASTA obtained from ThermoFisher (https://assets.thermofisher.com/TFS-Assets/LSG/manuals/ERCC92.zip). To distinguish reads from each transcriptome, we treated orthologous transcripts from each of the two frog species and *X. laevis* short (S) and long (L) subgenomes as independent ‘isoforms’. We therefore used kallisto RNA-seq software (v0.44.0; parameters: --bias --fr-stranded -t 4 --pseudobam) ^52^ to pseudo-align k-mers generated from reads to transcript sequences in a strand-specific manner. This approach can more accurately quantify ‘isoforms’ arising from the same transcript ^52^ and with our *Xenopus* hybrid transcriptome we can avoid complications from substantial multi-mapping or erroneously mapping to the wrong subgenome. We find that the majority of genes detected in the early stages of the hybrid are *X*. *laevis* maternal genes, as expected, and that *X. tropicalis* zygotic gene expression increases gradually through the first few time points and then more rapidly at the MBT (Figure S3A). Furthermore, we could accurately distinguish highly expressed maternal *X. laevis* short (S) and long (L) subgenome transcripts from their cognate *X. tropicalis* zygotic transcripts for the maternal mRNA of several key transcription factors, including *vegt*, *sox3*, *pou5f3.3*, and *ets2* ^21^ (Figure S3E).

### Normalization with exogenous spike-in RNA

The transcriptome composition across early embryogenesis is extremely dynamic compared to somatic cells. Maternal mRNAs are degraded while the number of expressed zygotic genes increases from zero to thousands of genes in several hours. Therefore, standard RNA-seq normalization approaches using the background transcriptome or library size are inappropriate. We therefore use absolute normalization with exogenous spike-in RNAs added during embryo collection to normalize *Xenopus* transcripts across time points. ERCC Spike-ins showed reproducibility across replicates and time points, and matched their expected abundance (Figure S2A-C). This resulted in smooth gene expression curves for many canonical zygotic genes and allows us to perform rigorous quantitative analysis on the resulting data. Transcript counts in each sample were normalized to the 38 most highly expressed ERCC spike-in RNAs by first calculating the geometric mean of all values for each ERCC gene, dividing an ERCC RNA’s geometric mean by each ERCC value at each time point, and taking one geometric mean of all values per condition-timepoint sample. This ‘scaling-factor’ is then used to multiply each gene expression value (raw read counts per transcript) at that time point. For example, all 10 hybrid and 10 cybrid samples were adjusted by the same size factors within replicate 2. This allows each replicate to be a self-contained normalized experiment. ERCC spike in transcript counts were compared to expected ratios and found to exhibit excellent reproducibility across replicates and experimental conditions. Two samples (replicate 2, cybrid 8.5 hpf; replicate 3, hybrid 7.0 hpf) had extreme technical variance, and these and their time-matched samples in the other treatment group (rep 2, hybrid 8.5 hpf; rep 3, cybrid 7.0) were removed from the ERCC normalization and all subsequent analysis. For the remaining high quality samples, ‘normalized expression’ in the main text refers to ERCC spike-in normalized expression and these counts were used in all analyses unless otherwise noted.

### Post-processing and filtering of transcripts

Prior to TPM calculations, we removed several genes with extremely high read counts due to homology with rRNA, or mitochondrial RNA, whose reads were not removed during the filter step (*Xetrov90027392, Xetrov90027395, dnajc28_1, Xelaev18000045m, Xelaev18003735m, Xelaev18003967m, slc41a2_1, Xelaev18002981m.g, LOC108645507*). We also removed all tRNA entries for similar reasons.

We found using BLAST and manual curation that the earliest and most highly expressed zygotic transcript, miR-427, had multiple FASTA entries in version 9.1 of the *X. tropicalis* transcriptome. Some entries were exact duplications and others were transcript variants. Because miR-427 is transcribed from repetitive loci and its transcript variants are difficult to quantify, we combined all read counts from these loci into a single entry, named “*Xetrov90009984m*” in our data set. The following miR-427 loci counts were combined: *LOC108646208, LOC108646207, LOC108646206, LOC101732110, LOC108646205, LOC108646204, LOC108646201, LOC108646202, LOC108646203, Xetrov90009976m, Xetrov90009977m, Xetrov90009978m, Xetrov90009979m, Xetrov90009980m, Xetrov90009981m, Xetrov90009982m, Xetrov90009983m, Xetrov90009984m, Xetrov90009985m, Xetrov90009986m, Xetrov90009987m, Xetrov90009988m, Xetrov90009989m, Xetrov90009990m, Xetrov90009991m, Xetrov90009992m, Xetrov90009993m, Xetrov90009994m, Xetrov90009995m, Xetrov90009996m*.

After the above filtering procedures, we re-calculated estimated TPM values per gene from ERCC spike-in normalized kallisto-derived counts using the formula: 10^6^ * ((estimated_counts / effective_length) / (SUM (estimated_counts / effective_length))). These TPM values were used in the determination of our ZGA gene set. Count values were then used for all other analysis unless noted otherwise. We used spike-in normalized count values because counts are more directly comparable across conditions and time points than TPM.

### Sequencing read alignment to genomes

To detect intronic RNA expression and generate read pileup tracks over the genome (Figure 2D; S3C) we first generated a hybrid genome FASTA file that contained the v9.1 *X. laevis* and v9.1 *X. tropicalis* genomes and the ERCC spike-in RNA transcriptome, and a corresponding GTF annotation file. This ‘xla_xtr_ercc’ genome was indexed using STAR v2.7.1a ^53^ (parameters: --sjdbOverhang --genomeChrBinNbits 15). rRNA, tRNA, and mitochondrial RNA filtered paired reads were aligned to the genome in a transcriptome-guided manner using STAR (parameters: --outFilterMultimapNmax 1 --alignEndsProtrude 10 ConcordantPair --outSAMmultNmax 5 --alignIntronMax 300000 --alignMatesGapMax 300000 --alignSJoverhangMin 8 --alignSJDBoverhangMin 1). Multi-mapping reads were removed and resulting bedGraph files from common time points in replicates 2 and 3 were merged. BigWig files were generated using kentutils bedGraphtoBigWig and signal tracks visualized with IGV (Broad Institute).

### Identification of zygotic transcripts

To analyze expression dynamics in embryos of different cellular DNA content, we first needed to define the set of *X. tropicalis* ZGA genes. In principle, every sequencing read mapped to *X. tropicalis* should be from a zygotically expressed transcript. However, the high maternal expression of highly similar *X. laevis* mRNA sequences may allow a low level of cross-contamination due to kallisto using a 31-mer k-mer based approach to assign reads to transcripts. Thus, even if only 5% of the reads from a highly expressed *X. laevis* maternal mRNA were mis-aligned to *X. tropicalis*, this could result in false positive expression and mis-characterization of that gene as zygotic *X. tropicalis*. In addition, the expression profile of many genes with low blastula or MBT expression that express highly in gastrulation could present as noisy data and are not useful for time-series comparisons. Therefore, we applied filters to *X. tropicalis* genes to identify a conservative, but high-confidence zygotic gene set. We first selected for *X. tropicalis* genes with expression less than 6 TPM (∼30-90 raw estimated counts, depending on gene length) in the first two time points, to remove genes with any potential contamination from *X. laevis* maternal reads. Next, we selected genes whose expression increased over the time series at least 8-fold between the sum of the first two time points and the sum of the 9.0 and 9.5 hpf time points.

We applied these thresholds to the hybrid and cybrid expression data from replicate 2 and replicate 3, which resulted in ∼1200-1400 ‘ZGA’ genes per condition. We retained 722 genes common in all 4 data sets to allow comparison between hybrids and cybrids. We then removed genes with fewer than 10 counts total in time points 9.0+9.5 hpf, removed provisionally annotated histone genes due to their repetitive loci origin, and summed counts for 5 genes that had 17 duplicate FASTA sequences with erroneous unique IDs. To further remove any possibility of cross-alignment contamination from *X. laevis*, we used BLAST to assess homology between the two frog species’ transcriptomes. We removed 22 genes with a contiguous region of 97% sequence identity over 100 nucleotides that was greater than 30% of the *X. tropicalis* gene’s mRNA length (*LOC100485134, LOC100486150, LOC100486870, LOC100490072, LOC100495041, LOC100496181, LOC100497915, LOC100498329, LOC100498553, LOC101732029, LOC108648872, actb, bin1, cbx1, cenpo, h2bc12, hes5.8, sox17a, sox18, sox18_1, sox18_2, tspan36*). The remaining 595 genes were used as the ‘ZGA gene set’ in all subsequent analyses unless otherwise noted.

### Fold-change calculations

We reasoned that a subtle shift in time-series expression between hybrid and cybrid conditions might not be detected if replicate expression values were merged initially because the variability in MBT timing between different clutches can be as high as 1-2 cell cycles (∼30-60 min). To assess differences in time-series between hybrids and cybrids, we therefore determined the log2-fold-change at each time point per replicate, using the hybrid as a reference, such that a negative fold-change indicates less expression in the cybrid. For plots in Figure 3, replicate log2-fold-changes were averaged across the 2-3 replicates at each time point.

### Gene activation time detection algorithm

We reasoned that any change in the activation time between hybrids and cybrids may represent a shift in transcription dynamics near the MBT, and therefore devised an algorithm to rigorously estimate the “activation time” for each gene. We note that this activation time does not correspond to the first transcription event in the embryo for a given gene (e.g., we detect expression earlier than the activation time using qPCR; data not shown). Rather, activation time is the time of large-scale transcript level increase from background cleavage-stage transcription levels to the level that occurs near the MBT. Thus, our activation time likely represents a “boost” in total embryonic transcription levels for a gene. This could be due to an increase in transcription rate at each promoter, or an increase in the number of cells expressing the transcript (at a constant rate), or both. While such an activation time could be obtained from the observed time series using a manual threshold, this threshold would depend on the observer so that we would not be confident in reporting differences in the 30-60 minute time scale required for our analysis.

In order to identify activation times that occur between data points, we used our data to generate a continuous estimate of gene expression. To do this, we fit the observed time series data to a cubic smoothing spline using SciPy’s Univariate Spline function (SciPy v1.5.2 Reference Guide; https://docs.scipy.org/doc/scipy/reference/generated/scipy.interpolate.UnivariateSpline.html). Smoothing splines are well-suited for interpolation of RNA-Seq data for two reasons. First, unlike exponential or polynomial regressions, smoothing spline regression does not require *a priori* assumptions about the shape of the underlying function. Second, in contrast to polynomial splines, a smoothing spline allows us to control how tightly the curves fit to (potentially random) variation in the data. The smoothing parameter p varies from 0 to 1 and determines the tightness of the fit. When p=0, the function passes through each data point, while for p=1, the function is a linear fit.

Prior to fitting the time series to the smoothing spline, it was first necessary to remove data points that were clear outliers because these outliers can drive estimates of the activation time. During the experiment, we expect gene expression to either remain constant (prior to zygotic gene activation) or monotonically increase (following activation). Therefore, we “flag” any upwards or downward ‘spike’ in the time series for potential removal. We consider the point to be an upward spike if the mean-normalized count value is more than 0.2 greater than both the previous and the subsequent time points. Similarly, a downward spike was taken to occur when the mean-normalized count value is more than 0.2 less than the previous and the subsequent time points (Figure S6). The above procedure may prove too stringent when a point is flagged due to its deviation from another spike and not from the trajectory of the remaining points. To avoid erroneously discarding a data point, we discard only one of every pair of adjacent flagged points according to the following procedure: First, each point is “unflagged” and fitted to a smoothing spline with p = 0.69, excluding the remaining adjacent point. Next, the point with the lowest distance to the resulting curve, is retained. Since this procedure requires fitting non-flagged points to a cubic smoothing spline, we discard any time series with greater than four flagged points before resolving adjacent flags. Of 595 original genes, 550 were retained across all four conditions (2 replicates, hybrid and cybrid). We note that special consideration must also be given to the first and last points in the time series, which cannot be flagged as spikes by virtue of lacking a previous or subsequent point, respectively. While early expressed time points did not show significant deviations from constant expression, the last point was occasionally significantly lower than the next-to-last point. In the latter case, we flag both the last and next-to-last points and discard one according to the general procedure for resolving adjacent flags described above.

We define the activation time as the first time point in the interpolated smoothing spline when the observed count level exceeds 20% of the spline’s maximum value on the interval 5-10 hours. While other threshold cutoffs and metrics based on the first or second derivative of the spline fit were possible ^59^), this method best accorded with visual inspection of gene curves. Our estimate of activation time will have error associated with the experiment and with the algorithm. We therefore sought to identify the smoothing parameter that minimized the error associated with the algorithm, which corresponds to minimizing the variation from experiment to experiment. As a way of doing this, we selected the smoothing parameter that maximized the inter-replicate activation time Pearson correlation coefficient for the top 100-highly expressed genes. The resulting optimum parameter was 0.69. We determined activation times for 547 of the 550 genes retained post-filter.

### Analysis of transcription dynamics per genome

Although we defined activation time from the time course of RNA-Seq count data, much of this increase likely results from the exponentially increasing number of genomes. Even in the absence of transcriptional activation at a single gene locus, a constantly expressed gene would still show activation-like kinetics owing to the exponential increase in number of genomes during the rapid embryonic cell division cycles.

To account for the potentially confounding effect of genome copies on our estimates of activation time, we sought to analyze the time course of transcripts per genome. We inferred a continuous function for genome equivalents (cell count) over time by defining a piecewise exponential function with doubling time between each observed cycle corresponding to the experimentally observed cell cycle period. The fact that this function is continuous allows us to approximate the increase in genomic material that occurs within S-phase of a single cell cycle. Transcripts per genome were then determined by dividing each count value by the inferred number of genomes at each time point. Unfortunately, normalizing by exponentially increasing numbers of genomes increased the prevalence of noisy fluctuations at low-expression values. Many of the resulting transcripts per genome curves showed an initial decrease before activation, but this concavity was seldom consistent across replicates. Despite this caveat, we were still able to reliably identify activation times from the transcripts per genome time courses using a modified version of the algorithm described above for the total transcription time courses.

More specifically, we defined the activation time as the first interpolated time point where the spline fit attained greater than 20% of maximum expression and had both positive first and second derivatives. The resulting activation times are well-correlated with the activation times determined from the transcript count series but are on average ∼30 min earlier. This indicates that the relative ordering of transcriptional activation times remains consistent whether activation is defined on an embryo-wide or per genome basis.

## QUANTIFICATION AND STATISTICAL ANALYSIS

Embryo samples were collected in a non-blinded way for the hybrid and cybrid treatment conditions. Individual embryos from each treatment group were randomly selected for each time point. Tubes for RNA extraction, cDNA generation, and library prep steps were processed in time-series order, alternating between hybrid and cybrid conditions, to avoid time-point batch effects during sample processing. Data was analyzed using R, Python, and Unix. Statistical tests performed in each panel are described in the corresponding figure legends. Pearson correlation tests and two-sample Kolmogorov-Smirnov tests were performed in R (version 4.0.2). Paired sample t-tests, two-tailed t-tests with unequal variance, and additional Pearson correlation tests were performed using Python (version 3.8). When appropriate, Bonferroni’s correction was applied to tests for multiple comparisons. Statistical approaches related to spline parameter selection, fitting smoothing splines to the expression time-series data, and the determination of activation times is described in detail in the methods. Asterisks in figure panels indicate significant differences in mean values between comparison groups, with corresponding p-values listed in the figure legends.

